# Combining systems and synthetic biology for in vivo enzymology

**DOI:** 10.1101/2024.02.02.578620

**Authors:** Sara Castaño-Cerezo, Alexandre Chamas, Hanna Kulyk, Christian Treitz, Floriant Bellvert, Andreas Tholey, Virginie Galéote, Carole Camarasa, Stéphanie Heux, Luis F. Garcia-Alles, Pierre Millard, Gilles Truan

## Abstract

Enzymatic parameters are classically determined *in vitro*, under conditions that are far from those encountered in cells, casting doubt on their physiological relevance. We developed a generic approach combining tools from synthetic and systems biology to measure enzymatic parameters *in vivo*. In the context of a synthetic carotenoid pathway in *Saccharomyces cerevisiae*, we focused on a phytoene synthase and three phytoene desaturases, which are difficult to study *in vitro*. We designed, built, and analyzed a collection of yeast strains mimicking substantial variations in substrate concentration by strategically manipulating the expression of geranyl-geranyl pyrophosphate (GGPP) synthase. We successfully determined *in vivo* Michaelis-Menten parameters (*K*_M_, *V*_max_ and *k*_cat_) for GGPP-converting phytoene synthase from absolute metabolomics, fluxomics and proteomics data, highlighting differences between *in vivo* and *in vitro* parameters. Leveraging the versatility of the same set of strains, we then extracted enzymatic parameters for two of the three phytoene desaturases. Our approach demonstrates the feasibility of assessing enzymatic parameters directly *in vivo*, providing a novel perspective on the kinetic characteristics of enzymes in real cellular conditions.

## Introduction

Enzymes catalyze most of the chemical reactions in living systems. A comprehensive interpretation of enzymatic parameters is therefore of paramount importance to grasp the complexity of cellular metabolism. The vast amount of data collected for thousands of enzymes has contributed to significant progress in our understanding of the remarkable chemical capabilities of biocatalysts and of their roles in cellular reactions.

Enzymatic reactions are traditionally analyzed *in vitro*, under dilute conditions, using pure or semi-pure protein samples in buffer solution. In contrast, the cellular medium is most accurately viewed as a heterogeneous, dense, crowded gel, containing various types of macromolecules and cellular lipidic organelles, with potential partitioning effects and variations in substrate and/or product diffusion coefficients (1). The parameters determined in classical enzymology experiments may therefore not be representative of *in vivo* reaction rates and equilibrium constants (2, 3). While some progress has been made in implementing and understanding viscosity and crowding effects in *in vitro* enzymatic assays, these conditions do not mimic the intrinsic complexity of the cellular environment (2).

*in vivo* enzymology seems to be the obvious approach to measure enzymatic parameters inside cells. Early attempts were made using the enzyme photolyase, for which both *in vitro* and *in vivo* parameters were determined (4, 5). Meanwhile, the *in vivo* V_max_ values of ten central carbon metabolism enzymes determined in the early 90s revealed significant differences between *in vivo* and *in vitro* assays for heteromeric protein complexes (6). The past ten years have seen growing interest from systems biology in determining the *in vivo* k_cat_ of native enzymes in model organisms. The *in vivo* apparent k_cat_ of E. coli enzymes have been determined independently by two research groups, leveraging advances in absolute protein quantification and high throughput metabolomics (7, 8). These values were obtained by dividing the flux of enzymatic reactions by the absolute abundance of the corresponding enzymes. Both studies found correlations between *in vivo* and *in vitro* data (with correlation coefficients around 0.6), suggesting that this method could serve as an alternative to *in vitro* assays. While this systems biology approach has provided valuable information, it cannot be employed to determine apparent *K*_M_s *in vivo*. Zotter et al. measured the activity and affinity of TEM1-β lactamase in mammalian cells *in vivo* with confocal microscopy, using a fluorescently tagged enzyme and a fluorescent substrate and product (9). They observed that the catalytic efficiency (*k_cat_/K_M_*) of this enzyme differs *in vitro* and *in vivo* due to substrate attenuation, indicating that *in vitro* data are not always indicative of *in vivo* function. Zotter et al. is probably the most comprehensive *in vivo* enzymology study to date, but this approach cannot be generalized because of a lack of universally appropriate fluorescent substrates/products for all enzymes. In a recent investigation of the *in vivo* kinetic parameters of thymidylate kinase (TmK), an interesting finding was the difference in TmK’s activity pattern when the substrate (thymidine monophosphate, TMP) was supplied in the media versus provided by internal metabolism, with Michaelis-Menten kinetics in the former and Hill-like kinetics in the latter. The authors hypothesized that the limited diffusion of TMP might be due to its confinement in a putative metabolon in E. coli (10).

*in vivo* enzymology is a particularly attractive prospect for membrane and multimeric proteins (6), which are tedious to purify and for which activity assays are difficult to optimize, mostly because artificial membranes are required (11–15). In the industrially important carotenoid pathway for example, despite the expression of numerous carotene biosynthesis enzymes, our understanding of their enzymatic behavior remains limited because many are membrane-associated proteins. While kinetic parameters for some phytoene synthases, sourced from plants or bacteria, have been established *in vitro*, many enzymatic assays require the co-expression of geranylgeranyl pyrophosphate (GGPP) synthase to attain activity, possibly because of the amphiphilic nature of the substrate or the requirement of membranes (12, 15). Another example of the difficulty of *in vitro* assays is phytoene desaturase, for which discernible enzyme activity has only been achieved in engineered environments (e.g. liposomes) with cell-purified substrate (14, 16–18).

This study combines synthetic and systems biology tools to develop an original *in vivo* enzymology approach. Synthetic biology offers a remarkable set of genetic engineering tools to precisely modulate the activity of enzymes in synthetic pathways, enabling *in vivo* control of substrate concentrations for the studied enzymatic reactions. In turn, systems biology can provide quantitative data on these reactions (fluxes and enzyme, substrate and product concentrations) and computational tools to model their behavior. We applied this approach to investigate a synthetic carotenoid production pathway in *Saccharomyces cerevisiae*, with industrial applications ranging from foods to pharmaceuticals.

## Results

### General principle of the proposed *in vivo* enzymology method

Enzymes are usually characterized in terms of their affinity (K_M_) and activity (maximal reaction rate V_max_ and turnover number k_cat_) (Fig 1). These parameters are typically determined *in vitro* by varying the substrate concentration across a relatively broad range (about two orders of magnitude) and measuring the reaction rate for each substrate concentration (19). Different mathematical formulas, such as the Michaelis-Menten equation, have been derived to estimate enzymatic parameters from these data. We propose a novel approach wherein the substrate concentration is varied directly within cells (Fig 1), by modulating the concentration of the enzyme producing the substrate of the studied reaction. Levels of the substrate-producing enzyme are varied using different combinations of promoter strengths and gene copy numbers, and translate into a wide range of substrate concentrations (2-3 orders of magnitude, as detailed in the following sections). In our experimental setup, each substrate concentration is achieved using a specifically engineered yeast strain. In each of these strains, the gene encoding the enzyme of interest is expressed at a given level, mirroring the conditions of *in vitro* experiments. Engineered strains are then grown under steady-state conditions (i.e., exponential growth), where reaction fluxes and metabolite concentrations remain constant (20). This stability allows for the accurate measurement of product formation fluxes (equivalent to reaction rates), intracellular substrate concentrations, and enzyme concentrations. The data obtained can then be combined to calculate apparent *in vivo* kinetics parameters, denoted K^cell^_1/2_, V^cell^_max_ and k^cell^_cat_ in analogy with the classical K_M_, V_max_ and k_cat_ parameters (Fig 1).

**Figure 1.**
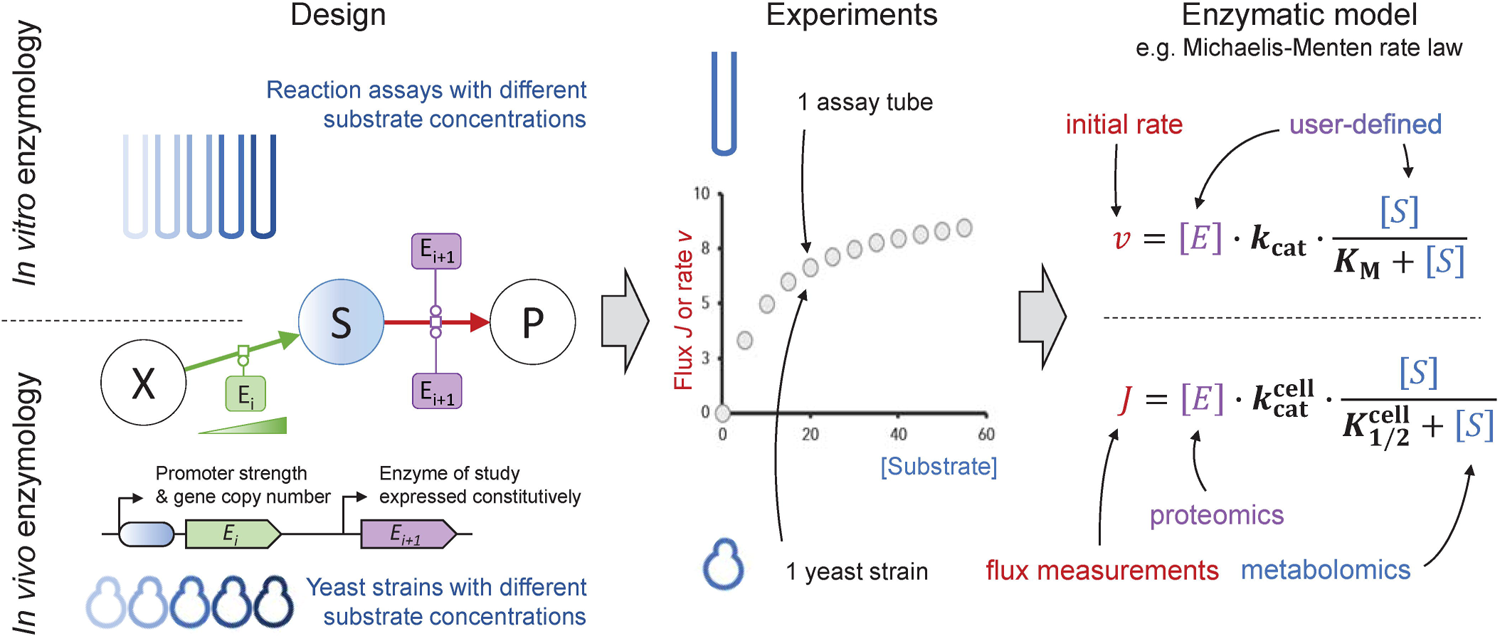
General principle of the proposed strategy for *in vivo* enzymology and comparison between *in vitro* and *in vivo* enzymology approaches.

### Construction of yeast strains to investigate a synthetic carotenoid production pathway

We evaluated our strategy by investigating a synthetic carotenoid pathway in yeast (*S. cerevisiae*), an industrially important chassis in biotechnology, which includes two membrane-interacting enzymes that are challenging to investigate *in vitro* (12, 16, 17, 21): phytoene synthase (CrtB) and phytoene desaturase (CrtI). Phytoene synthase, the first enzyme in the carotenoid pathway and considered the bottleneck of carotenoid biosynthesis in plants (21), condenses two molecules of geranylgeranyl pyrophosphate (GGPP) head-to-head to form phytoene. Phytoene synthases have proved difficult to express in soluble and active forms in microorganisms (12, 15, 21). *in vitro* rates and affinity constants have been determined for both plant and bacterial phytoene synthases but these parameters may have been altered by the conditions of the *in vitro* assays—adding detergents for the bacterial enzyme and co-expressing a GGPP synthase or using semi-crude extracts for the plant enzyme (12, 15, 21–24). Following this enzymatic step, three insaturations are introduced in phytoene to produce lycopene. In non-photosynthetic bacteria and fungi, three reaction steps are catalyzed by a single enzyme (CrtI in bacteria and CarB in fungi) (25), whereas in plants and cyanobacteria, four different enzymes are involved in the conversion of phytoene to lycopene (26). Given that these desaturation steps are a major bottleneck in microbial carotenoid biosynthesis, determining *in vivo* enzymatic constants is an industrially relevant challenge (10, 22, 23).

To analyze these enzymes, we assembled a set of strains covering a broad range of substrate concentrations (Fig 2 and Table EV1). To create the collection of strains with different intracellular levels of GGPP, the substrate of phytoene synthase, we first increased GGPP production by modifying the native terpene pathway: strong constitutional expression of *ERG20* and of the truncated form of *HMG1* and deletion of the main phosphatase gene involved in GGPP and FPP pyrophosphate group removal (*DPP1*), yielding the strain yENZ15 (29, 30). Three different integration cassettes containing different microbial GGPP synthases (CrtE) expressed under different constitutive promoters were integrated into the yENZ15 genome, yielding a series of strains designed to produce a wide range of GGPP concentrations (Fig 2).

**Figure 2.**
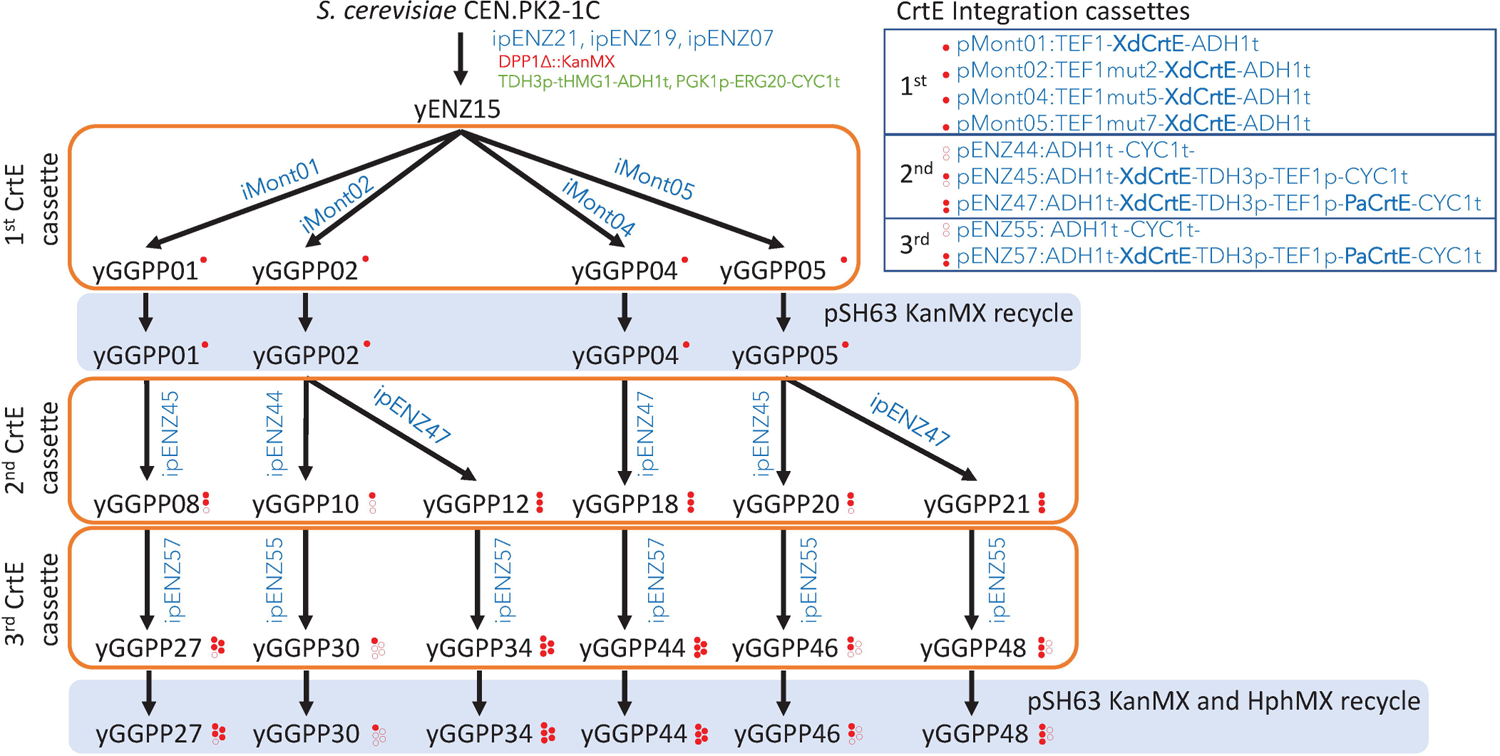
Scheme of the construction of yeast strains with different GGPP concentration. Red filled dots represent a copy of GGPP synthase, and empty dots represent an empty integration cassette.

We then verified that the intracellular GGPP concentration was indeed modulated *in vivo*, as done previously (24, 32–34). GGPP concentration varied by a factor of 38 in these strains (Fig 3B), and the strains all had similar growth rates (0.40±0.03 h^-1^) (Fig 3B) indicating that there was no change in cell physiology. As detailed below, we used this set of strains as a base to study a phytoene synthase (https://www.uniprot.org/uniprotkb/P21683/entry, EC 2.5.1.32) and three different lycopene-forming phytoene desaturases (EC 1.3.99.31).

**Figure 3.**
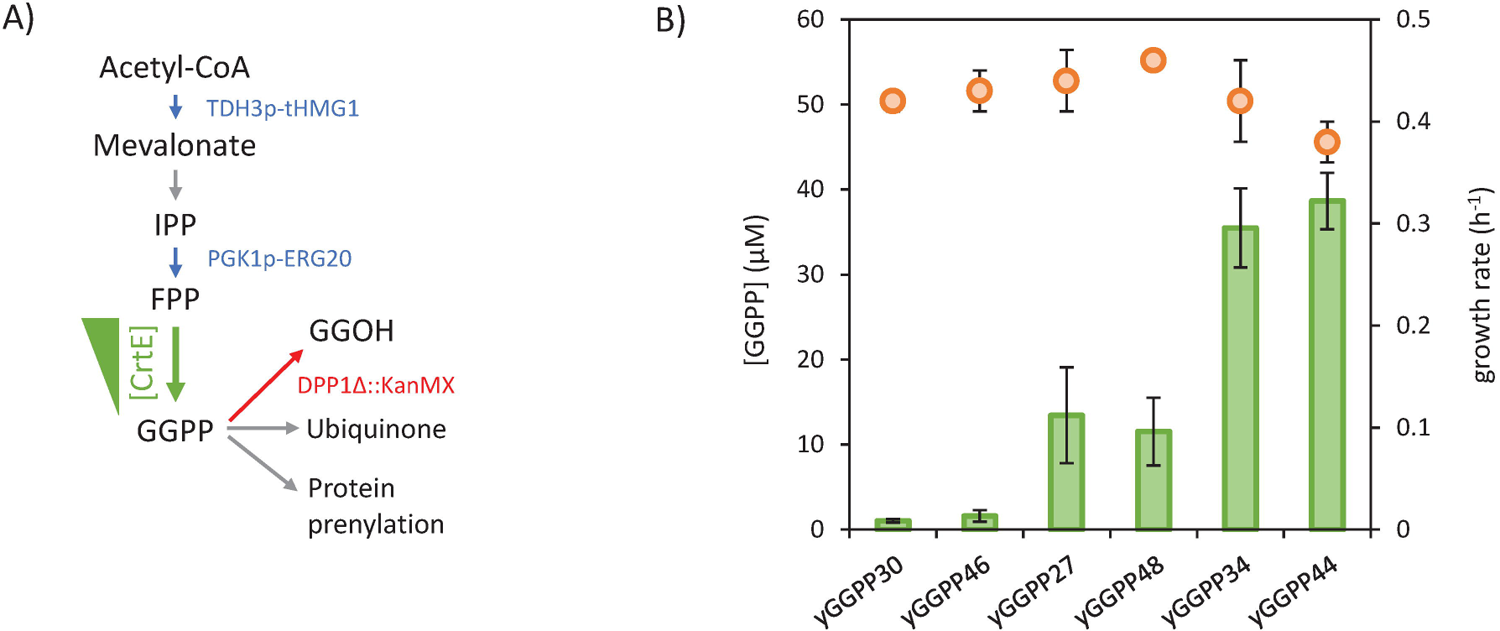
Construction of the set of yeast strains with different intracellular GGPP concentrations. A Genomic modification scheme for the yENZ15 strain and intracellular fates of GGPP in *S. cerevisiae*. Overexpressed enzymes are shown in blue, enzyme deletion is shown in red, and enzyme modulation is shown in green. B GGPP concentration (green bars) and specific growth rate of *S. cerevisiae* strains (orange dots). Mean values ± standard deviations (error bars) were estimated from three independent biological replicates.

### *In vivo* enzymatic parameters of phytoene synthase from Pantoea ananas

We used the set of strains with different intracellular levels of GGPP (Fig 3) to study phytoene synthase from *Pantoea ananas* (PaCrtB) (Fig 4A). In a classical enzymatic reaction setup, enzyme concentration must be carefully controlled to establish an informative set of reaction rate vs substrate concentration data points. To evaluate the impact of enzyme concentration on the collected kinetic profiles, we developed a kinetic model and performed simulations for three different enzyme concentrations (low, medium and high) under a broad range of substrate-producing flux. The relationships between the steady-state substrate concentration and the flux are shown in Fig 4B. Simulation results suggest that excessively high enzyme concentrations would hinder substrate saturation, rendering parameter estimation impossible. In contrast, an extremely low enzyme concentration would lead to a low production flux, making precise measurements difficult and reducing the chance of obtaining accurate enzyme parameters. Therefore, we decided to perform our saturation experiments using three enzyme concentrations by expressing *P. ananas crtB* under the control of three different constitutive promoters (low expression: TEF1mut2p, medium expression: PGI1p and high expression: PDC1p).

**Figure 4.**
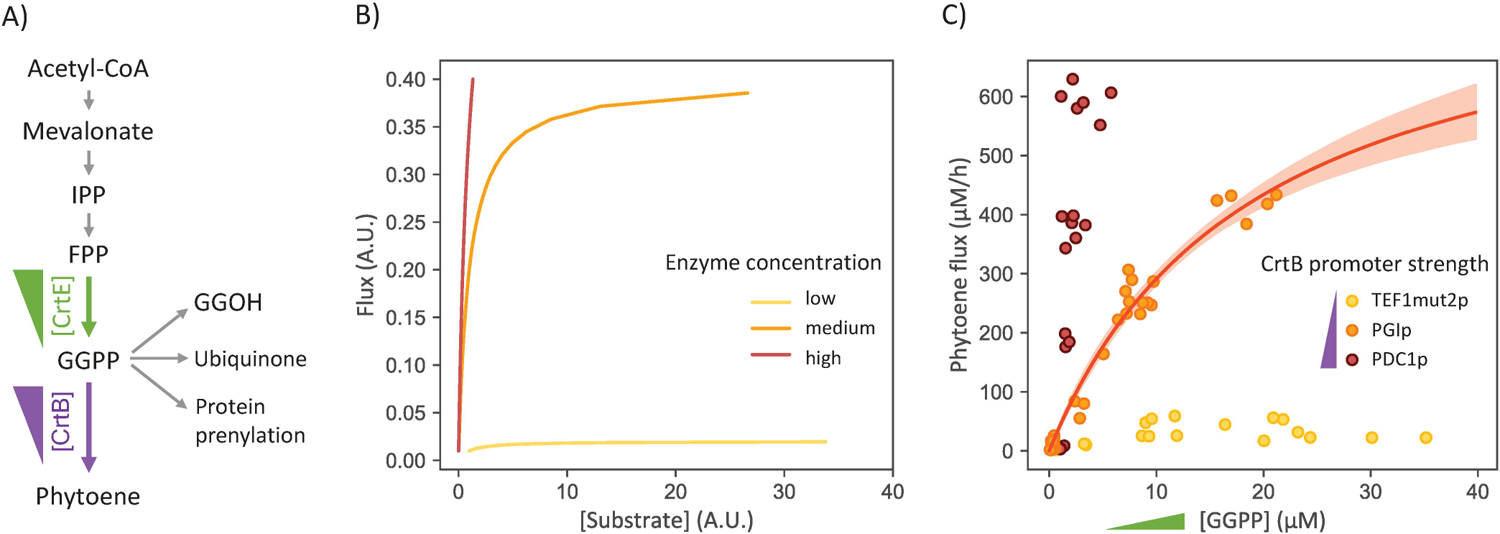
*in vivo* characterization of phytoene synthase from Pantoea ananas. A Scheme for the biosynthesis of phytoene by CrtB. B Steady-state flux and substrate concentration simulated for three different CrtB activities (low, medium and high) under a broad range of GGPP-producing flux. C *in vivo* characterization of CrtB in *S. cerevisiae*: phytoene production as a function of GGPP concentration in strains with low (TEF1mut2p), medium (PGIp) and high (PDC1p) level of CrtB. Each data point represents an independent biological replicate. For PGIp strains, red line represents the best fit of a Michaelis-Menten rate law, and shaded area corresponds to 95 % confidence interval on the fit.

Consistent with simulation results (Fig 4B), the GGPP concentration progressively decreased as PaCrtB activity increased (Fig 4C). Therefore, the GGPP concentration was very low at high PaCrtB level (PDC1p), and enzyme saturation was not reached. In contrast, the phytoene flux increased as PaCrtB expression increased. The phytoene concentration was too low for accurate quantification of the phytoene flux at low levels of PaCrtB (TEF1mut2p). However, accurate measurements of GGPP concentration and phytoene flux could be made at medium PaCrtB levels (PGIp). Based on these results, we used strains with medium PaCrtB levels for further investigations. A 167-fold variation in GGPP concentration (from 0.12 to 20.78 µM) was observed in these strains. Partial saturation was achieved, leading to a nonlinear relationship between the phytoene flux and the GGPP concentration, as observed in classical *in vitro* experiments. We assumed and verified, by western blot and mass spectrometry-based proteomics, that PaCrtB levels were identical in all strains (Fig EV1 and Table EV2), indicating that PaCrtB activity (V^cell^_max_) was also similar in all strains. Precise estimates of V^cell^_max_ (849±71 µM/h, rsd = 13 %) and K^cell^_1/2_ (19±3 µM, rsd = 14 %) were obtained by fitting these data using an irreversible Michaelis-Menten model (consistent with previous reports (21) and with the very negative ΔG^0^ value of this reaction) (Table EV3). As expected from the simulations, parameters could not be determined precisely from the two other datasets (Table EV3). The affinity of PaCrtB for GGPP estimated from our *in vivo* data is higher than measured *in vitro* (K_M_ = 41 µM) (21), highlighting the necessity of measuring enzyme parameters directly within the intracellular environment. Meanwhile, the k^cell^ of 6±1 s^-1^, obtained from absolute quantitative proteomics measurements of the enzyme concentration (42 nM) in strains expressing PaCrtB under the control of PGI1p, is comparable to the one obtained *in vitro* for CrtB in Pantoea agglomerans (14 s^-1^, Table EV4) (15).

The theoretical maximum phytoene production flux in these strains is therefore 849±71 µM/h. The bottleneck for carotenoid biosynthesis in plants is phytoene synthase, whose activity is regulated by a combination of transcriptional, post-transcriptional, and post-translational mechanisms to adjust carotenoid production (35). In our synthetic system however, our results suggest that GGPP biosynthesis is the limiting step since five copies of the GGPP synthase gene are required to achieve just 51 % of the maximum phytoene production flux, due to the lack of phytoene synthase saturation.

### Straightforward extension to analyze downstream enzymes of the carotenoid pathway: phytoene desaturases

The most time-consuming tasks in the proposed *in vivo* enzymology protocol are the molecular cloning and strain engineering steps required to generate strains with different substrate concentrations. However, the strains used to investigate a given enzyme can be recycled to investigate downstream steps in the same metabolic pathway, significantly streamlining experiments, which become a simple plug-and-measure process (Fig 5A). Integration of crtB, controlled by the high-expression promoter PDC1 into our initial set of strains, produced strains with a 430-fold variation in phytoene concentration (from 4 to 1641 µM). These strains thus form an ideal framework to investigate the *in vivo* kinetics of the next metabolic step with various phytoene desaturases.

**Figure 5.**
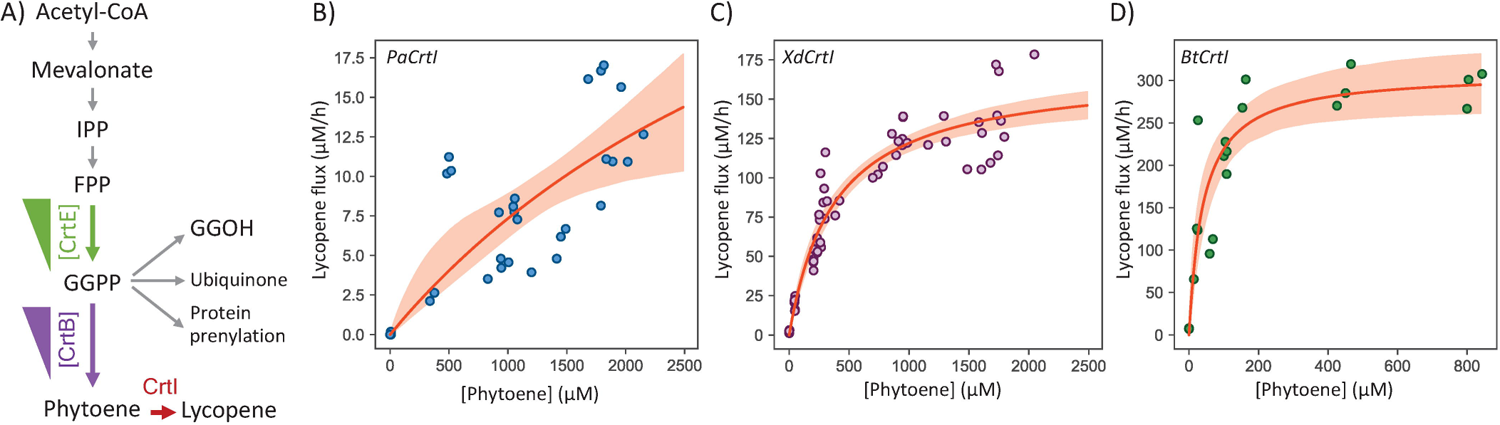
*in vivo* characterization of three phytoene desaturases. A Scheme for the biosynthesis of lycopene by CrtI. B–D *in vivo* characterization of Crtl from *P. ananas* (B), X. dendrorus (C) and B. trispora (D). Each data point represents an independent biological replicate, red lines represent the best fits of a Michaelis-Menten rate law, and shaded areas correspond to 95 % confidence intervals on the fits.

To demonstrate this point, we chose three widely used microbial lycopene-forming phytoene desaturases (1.3.99.31) (22, 23, 25, 26): *P. ananas* Crtl (P21685, PaCrtI), which, along with *Rubrivivax gelatinosus* Crtl, is one of only two lycopene-forming phytoene desaturases with published *in vitro* enzymatic parameters (14, 28); and two fungal CrtIs widely used in carotenoid production, one from *Phaffia rhodozyma*, formely *Xanthophyllomyces dendrorhous* (A0A0P0KMF9, XdCrtI), and the other from *Blakeslea trispora* (Q67GI0, BtCrtI), with no enzymatic data available to date.

As for the investigation of phytoene synthase, the expression of the phytoene desaturases was controlled by different promoters: TDH3p for PaCrtI, PGK1p for XdCrtI, and PGI1p for BtCrtI. These promoters were chosen based on the relative lycopene producing abilities of the different CrtIs in *S. cerevisiae*. This was especially important for BtCrtI, where high expression in the exponential phase caused cell death due to the formation of lycopene crystals (Fig EV2A). PaCrtI and XdCrtl levels did not vary with the phytoene concentration of the strains, but BtCrtI levels were slightly higher in the strains with the lowest phytoene concentrations (Fig EV3). The higher levels of PaCrtI compared with those of the fungal phytoene desaturases is explained in large part by the use of a TDH3 promoter. However, lycopene fluxes were 10 times lower with PaCrtl than with XdCrtl or BtCrtl (Fig 5). Phytoene levels were high enough to saturate both fungal CrtIs (Fig 5) but not PaCrtl, for which lycopene fluxes remained low for all tested phytoene concentrations.

The experimental data from the three enzymes were fitted with an irreversible Michaelis-Menten equation, from which V^cell^ and K^cell^ values were successfully obtained for XdCrtl and BtCrtl but not for PaCrtl, for which only lower limits for V^cell^ (13 µM·h^-1^) and K^cell^_max_ (560 µM) could be determined (Table EV3). The V^cell^_max_ values obtained for BtCrtI (309±22 µM·h^-1^) was higher than for XdCrtI (168±7 µM·h^-1^), and the K^cell^_max_ for BtCrtI (41±12 µM) was 9 times smaller than XdCrtI’s (380±49 µM). The k^cell^_1/2_ obtained by integrating quantitative proteomics measurements in strains expressing XdCrtI under PGK1p was 0.030±0.006 s^-1^ (Table EV3).

These results highlight the versatility of our approach to compare the *in vivo* behavior of enzymes from different organisms, using the same set of strains, and provide valuable information to understand and optimize natural and synthetic pathways *in vivo*.

## Discussion

Inspired by conventional *in vitro* enzyme assays, we developed an innovative approach to measure enzyme kinetics *in vivo*. We successfully used the proposed method to estimate the *in vivo* equivalents of Michaelis-Menten parameters for a phytoene synthase and two phytoene desaturases in *S. cerevisiae*.

These results highlight the reliability and versatility of our *in vivo* enzymology approach, which hinges on solving two methodological challenges: 1) controlling the substrate pool over a broad concentration range, and 2) measuring the total concentrations of substrate, product and enzyme. The first challenge was addressed using synthetic biology tools to modulate the concentration of a substrate-producing enzyme (here GGPP synthase, CrtE). The second challenge was overcome thanks to sensitive, quantitative metabolomics and proteomics techniques that can be generalized to various other enzymatic models. Indeed, the coverage of the metabolome and fluxome by current omics approaches is now high and is continuously increasing, which enables the application of the proposed approach to a broad range of enzymes. In this study, we quantified three basic enzymatic parameters (K^cell^_1/2_, k^cell^_cat_ and V^cell^_max_). In the absence of absolute quantitative proteomics data, the proposed method still allows for the determination of V^cell^_max_ and K^cell^_1/2_ values, providing sufficient information for most enzymology and metabolic engineering studies. Similarly, estimating the absolute K^cell^_1/2_ values can be achieved using only relative flux values, without requiring absolute concentration of enzymes.

We demonstrate that substrate concentrations can be varied *in vivo* over a wide range. Here, the GGPP concentration was varied by a factor 167 and the phytoene concentration by a factor 430, allowing clear enzyme saturation curves to be observed for the two fungal phytoene desaturases (XdCrtI and BtCrtI). However, while the range of substrate concentrations explored was significantly wider than those used in *in vitro* studies of the same enzymes (Table EV4) (16, 21), complete saturation was not achieved for PaCrtB or PaCrtI. It is conceivable that, when these enzymes are bound to natural membranes, their apparent affinity for the substrate could differ from the *in vitro* environment. A second explanation concerns substrate availability. While classical *in vitro* enzyme assays involve homogeneous, highly diluted buffers, the intracellular environment is dense, heterogeneous and compartmentalized. In this first *in vivo* approach, variations in substrate concentrations between compartments were not considered explicitly, though it may have affected the values obtained for the parameters. For example, a higher concentration of phytoene inside lipid droplets compared with the rest of the membrane would reduce phytoene concentrations around CrtI enzymes and would thus limit substrate saturation. Enzyme localization could also affect measurements as conditions (pH, concentration of ions, etc.) vary between compartments. Iwata-Reuyl et al. found for instance that the measured activity of PaCrtB is 2,000 times lower in the absence of detergents (15), and Fournié et Truan observed that different heterologous CrtI expression systems produced different phytoene saturation patterns (36). In the case of the two *P. ananas* enzymes, the mechanisms that cause the absence of saturation may be different. For PaCrtI, the observed concentrations of phytoene (substrate) and lycopene (product) are either equivalent to or lower than those obtained with the two fungal CrtI enzymes. This suggests a true difference in the affinity for phytoene between the bacterial and fungal enzymes. For PaCrtB, given that the GGPP concentration in the cell is high (around 20 µM), the lack of complete saturation is more likely due to a heterogeneous distribution of GGPP within the cell, with only a fraction of GGPP being in the vicinity of the enzyme. Dedicated studies will be required to clarify the effects of cell compartments on enzyme efficiency.

In addition, other explanations for the difficulty in reaching complete saturation merit attention as they also convey general hypotheses on metabolic systems: i) substrate or product toxicity detrimental to cell growth (e.g. the specific growth rate is nearly 50 % lower for lycopene concentrations above 400 µM, (Fig EV2B), ii) overflow mechanisms that relocate some of the substrate to a different cell compartment or even outside the cell (37, 38), iii) the presence of alternative pathways that divert additional substrate produced above a certain concentration threshold (39, 40), or iv) intrinsic properties of metabolic systems whereby production fluxes tend to decrease when product concentrations increase (41–43). While these mechanisms contribute to global metabolite homeostasis (44–46) and are essential for cell viability, they limit substrate accumulation and thus could prevent complete saturation of the enzyme. Moreover, it is crucial to bear in mind that, similarly to *in vitro* data, enzymatic parameters are only valid under the conditions used to measure them. We therefore argue that enzymatic parameters should not be considered constants since they depend on the microenvironment (pH, ion concentration, temperature, membrane composition, etc.), be that *in vitro* or *in vivo*. Still, as mentioned below, these parameters are useful for pathway engineering and an advantage of our *in vivo* enzymology method is that kinetics parameters are measured under the exact same conditions as the enzyme is expressed.

Our approach enables the determination of whether a given enzyme is operating at saturation *in vivo* under different cellular conditions. The tested enzymes displayed various saturation profiles, indicating that they may function under diverse conditions within the cell. Operating at full saturation (far above the K^cell^, the enzyme works at its maximum rate, being unaffected by changes in substrate concentration or minor environmental variations. This makes the enzymatic reaction highly robust in terms of product formation. Conversely, if an enzyme operates at substrate concentrations much lower than the K^cell^, any variation in substrate concentration will directly affect the production rate. This could maintain substrate homeostasis, potentially prevent toxic effects due to concentration changes. From a metabolic engineering perspective, achieving the optimal balance between high product formation and metabolic homeostasis is crucial. This balance can be attained by adjusting the levels of both the substrate-forming enzyme and the target enzyme. This process needs to be repeated for each enzyme, potentially under different saturation regimes. In biotechnology, our *in vivo* enzymology approach may thus guide metabolic engineering strategies to ensure that the overall pathway maintains the desired balance between production and cellular homeostasis, thereby ensuring greater stability and robustness of the engineered microbial strains at maximal production flux.

The final step in our *in vivo* enzymology method involves fitting experimental data with a mathematical model of enzyme kinetics to obtain the corresponding parameters, the same approach as used *in vitro*. Traditional enzymatic models were derived with *in vitro* measurements and assumptions in mind. For instance, the classical Michaelis-Menten relationship applies only at steady-state, a condition clearly met in our experimental setup where all data were collected during exponential growth (i.e. metabolite concentrations and fluxes remain constant over time). However, other assumptions do not hold *in vivo*, in particular regarding the product concentration, which cannot be zero. Nevertheless, for the enzymes investigated here, where the catalyzed reactions are essentially irreversible, the reverse reaction can be neglected, and the Michaelis-Menten formalism still applies. Another important assumption is that the substrate concentration must be higher than the enzyme concentration. In this study, the *in vivo* substrate/enzyme ratio was between 8 and 1400 for PaCrtB and between 2 and 2500 for XdCrtI (Table EV5). Surprisingly, this criterion was not met for CrtB and CrtI in previous *in vitro* assays (19, 37), with substrate/enzyme ratios between 0.05 and 0.18 for PaCrtB and between 1 and 2.57 for PaCrtI (Table EV4). In our setup, despite some formal assumptions not being fully satisfied, the data were still satisfactorily fit with a Michaelis-Menten equation. This underscores the applicability of our method, from which informative parameters such as the degree of saturation can be inferred to understand *in vivo* enzyme function and to engineer natural and synthetic pathways for biotechnology. The availability of *in vivo* data about enzyme kinetics may also lead to the derivation of specific laws that account for the specificities of *in vivo* studies (e.g., non-negligible product concentrations).

As we advocate for the simplicity of our method, we would like to share some insights on its implementation in future studies. First, to maximize the output of genetic constructs, experimental design should be employed to compare various enzymes that catalyze the same reaction (whether mutants or from different species) or to target multiple enzymes in the same metabolic pathway, in this case phytoene synthase and phytoene desaturase. Second, the number of genetic constructs necessary to reach saturation (ideally 4 to 5) can be minimized by verifying the substrate production range early on. Third, in most cases, testing three enzyme concentrations has been sufficient to obtain a satisfactory saturation curve or at least to precisely estimate the K^cell^ range. Lastly, improvements can be made using high-throughput methods for the construction of plasmids and strains, for growth experiments and for samples preparation steps. The development of single cell metabolomics and proteomics approaches will also significantly increase the throughput of the strain characterization step in future studies.

Sixteen years ago, in their review on enzyme function, Dagmar Ringe and Gregory Petsko wrote: “How do enzymes function in a crowded medium of low water activity, where there may be no such thing as a freely diffusing, isolated protein molecule? *in vivo* enzymology is the logical next step along the road that Phillips, Koshland, and their predecessors and successors have traveled so brilliantly so far” (3). The work presented here, albeit performed in the context of a synthetic metabolic pathway, touches on the difference between kinetic parameters measured *in vitro* and *in vivo* and their interpretation. We notably show how fine-tuning and balancing the expression of the substrate-producing enzyme and the enzyme under study yields datasets from which meaningful and reliable enzymatic parameters (K^cell^, k^cell^ and V^cell^) can be obtained. By including additional controlled steps, this method could be applied to a wider range of variables, such as inhibition and activation parameters (by modulating the pool of regulatory metabolites). This method could also benefit from dynamic data (time-course monitoring in response to metabolic or genetic perturbations). New formalisms will be required to account for *in vivo* conditions, notably the presence of products, similar enzyme and substrate concentrations, local concentration variations, and molecular fluxes within the cell. Our method is particularly valuable for studying membrane-bound and multimeric enzymes, for which the purification and assay optimization steps of classical *in vitro* enzymology can be extremely challenging. For membrane-bound enzymes, *in vivo* enzymology offers a realistic environment devoid of detergents or other interferences, with natural membranes rather than liposomes. We sincerely hope that our work will stimulate further studies delving deeper into how enzymes function in their natural environment.

## Materials and methods

### Plasmid construction

Plasmids and primers are listed in Tables EV6-7. Plasmid sequences and annotations are provided in Dataset EV1. The primers were synthesized by IDT (Leuven, Belgium) and the sequences of PaCrtB, PaCrtI and BtCrtI were codon optimized for yeast and synthetized by Twist Bioscience (San Francisco, California). XdCrtE and XdCrtI from *Enterobacter agglomerans* were amplified from pMRI34-CrtE-Gal1-10-HMG1t, YEplac195 YB/I, and pAC-BETA respectively (47–49). Sequences of the mutated versions of the TEF1 promoter TEF1mut2p, TEF1mut5p and TEF1mut7p were obtained from Nevoigt et al. (50). Polymerase chain reaction (PCR) was performed using Phusion high fidelity polymerase and Phire Hot start II DNA polymerase (ThermoFisher Scientific, Lithuania). DNA fragments were purified using Monarch DNA Gel Extraction Kit from New England Biolabs. DNA fragments were annealed by isothermal assembly using NEBuilder HiFi assembly kit from New Englands Biolabs. Clones and plasmids were propagated in homemade calcium- and TOP10-competent Escherichia coli cells.

### Construction of yeast strains

All yeast strains used in this study are derived from CEN.PK2-1C and are listed in Table EV1. Yeasts were transformed using Gietz et al.’s high-efficiency transformation protocol (51). Integrative cassettes were obtained by enzyme digestion or PCR and were used without any further purification. Strains were selected using auxotrophy markers or antibiotic resistance at a concentration of 400 µg/mL. Antibiotic resistance recycling was performed using vector pSH63 as described in the literature (52). Genome integration was verified by colony PCR using the primers listed in Table EV7. Genomic DNA was extracted using DNA release from ThermoFisher Scientific.

### Media and culture conditions

All strains were grown in modified synthetic Verduyn media containing glucose (111 mM), NH_4_Cl (75 mM), KH_2_PO_4_ (22 mM), MgSO_4_ (0.4 mM) and CSM (ForMedium LTD, Hunstaton, England) at pH 5.0 (53). Sterilization was performed by filtration. Fresh colonies from selective plates were precultured in 350 µL complete synthetic medium at 28 °C for 8 hours and these cells were used to inoculate cultures with a 1:5 medium:flask proportion to an initial OD_600nm_ of 0.002, grown at 200 rpm at 28 °C.

### Carotene quantification

Samples (10 or 20 mL) of yeast culture were harvested with an OD_600nm_ of approximately 5, centrifuged, and washed with 1 mL of MilliQ water. Cell pellets were freeze-dried and stored at −80 °C until extracted. β-apocarotenal solution (40 µL, 50 µM) was added to the dried cells, and carotenes were extracted with glass beads and 500 μL of acetone in three 20 s rounds of agitation at 0.05 m/s with a FastPrep FP120 cell disruptor (ThermoFisher). The acetone phase was transferred to a new tube and the extraction was repeated twice. Acetone extracts were pooled, centrifuged, dried under nitrogen flux, and resuspended in acetone for HPLC analysis. Analyses were carried out on a Thermo Scientific Vanquish Focused UHPLC Plus system with DAD HL. Extract samples (5 μL) were injected into a YMC carotenoid column (100 × 2.0 mm and 3 μm particle size) equipped with a precolumn (100 × 2.0 mm and 3 μm particle size). The mobile phases used to separate and quantify phytoene, lycopene and β-apocarotenal from ergosterol and derivatives were mixtures of (A) methanol/water (95:5) and (B) dichloromethane. The flow was 0.25 mL/min with the following gradient: 0–0.1 min 5 % B, 0.1–0.5 min 20 % B, 0.5–2 min 60 % B, 2–5 min 80 % B, 5–8 min 80 % B and 8–11 min 5 % B. The absorbance from 210 to 600 nm was measured throughout the run with a data collection rate of 2 Hz and a response time of 2 s. The phytoene concentration was deduced from its absorbance at 286 nm and lycopene and β-apocarotenal concentrations from the absorbance at 478 nm. The reference wavelength (600 nm) was subtracted from each of the wavelengths used for metabolite quantification.

### Flux calculation

Phytoene and lycopene are produced and accumulate in the cells, and their pools are continuously diluted by cell growth. Assuming an absence of degradation or reutilization of these end-products by the cell, phytoene and lycopene production fluxes are balanced solely by their dilution fluxes in the exponential growth phase, where cells are at metabolic steady-state. Thus, phytoene and lycopene production fluxes were determined by multiplying their concentrations by the cell growth rate. This flux calculation method provides results consistent with those obtained by targeted ^13^C-fluxomics (54).

### GGPP quantification

GGPP was quantified as detailed previously (55). Briefly, 10 mL of broth was filtered through 0.45 μm Sartolon polyamide (Sartorius, Goettingen, Germany) and washed with 5 mL of fresh culture medium (without glucose). The filters were rapidly plunged into liquid nitrogen and then stored at −80 °C until extraction. Intracellular GGPP was extracted by incubating filters in closed glass tubes containing 5 mL of an isopropanol/H_2_0 NH_4_HCO_3_ 100 mM (50/50) mixture at 70 °C for 10 min. For absolute GGPP quantification, 50 μL of ^13^C internal standard were added to each extract. Cellular extracts were cooled on ice and sonicated for 1 min. Cell debris was removed by centrifugation (5000 g, 4 °C, 5 min). Supernatants were evaporated overnight (SC110A SpeedVac Plus, ThermoFisher, Waltham, MA, USA), resuspended in 200 μL of methanol:NH_4_OH 10 mM (7:3) at pH 9.5 and stored at −80 °C until analysis.

Analyses were carried out on a LC–MS platform composed of a Thermo Scientific Vanquish Focused UHPLC Plus system with DAD HL, coupled to a Thermo Scientific Q Exactive Plus hybrid quadrupole-Orbitrap mass spectrometer (ThermoFisher), as detailed previously (55). Calibration mixtures (prepared at concentrations from 0.08 nM to 10 μM) were used to construct calibration curves from which the absolute concentration of GGPP in the samples was determined.

### Western blot

Protein extracts were prepared as described by Zhang et al. (46). Briefly, 1.5 OD_600nm_ of pelleted cells were pre-treated with 100 µL of a 2 M lithium acetate cold solution, and left to stand for 5 min, followed by 51min centrifugation at 50001g, 41°C. The supernatant was discarded and 100 µL of a 0.4 M solution of NaOH was added. After gentle resuspension, and 5 min standing on ice, the samples were centrifuged for 51min at 41°C. After discarding supernatants, the pellets were vigorously vortexed with 60 µL of bromophenol blue loading dye solution with 5 % β-mercaptoethanol. After denaturation for 101min at 991°C, 51µL of each sample was loaded onto 10 % SDS page gel. Semi-dry transfer was performed on PVDF membrane (Merck Millipore, Darmstadt, Germany) using a Trans-Blot SD Cell BioRad apparatus (181V during 201min), and 5 % bovine milk in TBS as blocking agent. Incubations were performed with mouse anti-FLAG or mouse anti-V5 (ThermoFisher Scientific), and secondary anti-mouse IgG coupled with horseradish peroxidase (ThermoFisher Scientific), diluted according to manufacturer instructions. Proteins were detected by incubation with SuperSignal West Pico PLUS substrate (ThermoFisher Scientific).

### Proteomics

For cell disruption, 10^8^ cells were dissolved in 200 µL of lysis buffer (0.1 M NaOH, 2 % SDS, 2 % 2-mercaptoethanol, 0.05 M EDTA), heated at 90 °C for 10 min and neutralized with 5 µL of 4 M acetic acid. Glass beads were added and the samples were vortexed at 4 °C for 30 min. Cell debris was pelleted by centrifugation and 3000 g for 10 min and the protein concentration of the supernatant was determined using the Bradford assay. Protein aliquots (400 µg) were cleaned by methanol chloroform precipitation. The protein samples were dissolved in 5 µL of 6 M guanidinium chloride, 5 µL of 0.1 M dithiothreitol (DTT), and 100 µL of 50 mM TEAB 50 and diluted to a protein concentration of 2 µg/µL with 90 µL of water. Absolute protein quantification was performed using heavy isotope labeled tryptic peptides as internal standards. Protein lysate aliquots (20 µg) were spiked with a mixture of ten AQUA peptides, C-terminally labeled with heavy lysine or arginine (ThermoFisher Scientific) with concentrations of 50 fmol/µg, 10 fmol/µg, 5 fmol/µg and 0 fmol/µg. The samples were reduced by adding 2 µL of 0.1 M DTT and incubating at 60 °C for 1 h, alkylated with 2 µL of chloroacetamide 0.5 M at room temperature for 30 min, and digested with 0.5 µg of trypsin overnight. Digestion was stopped by adding 5 µL of 1 % TFA and the samples were cleaned using 100 µL C18 tips. The extracts were lyophilized, reconstituted in 20 µL of eluent A, and transferred to HPLC vials.

Samples were analyzed in triplicate using an UltiMate 3000 UHLC system coupled to a Q Exactive Plus mass spectrometer (ThermoFisher Scientific). Approximately 0.6 µg of peptides were loaded on a C18 precolumn (PepMap100, 5 μm, 300 Å, ThermoFisher Scientific) and separated on a PepMap RSLC C18 column (50 cm × 75 μm, 2 μm, 100 Å, ThermoFisher Scientific) with a 1.5 h gradient with eluent A (Water, 0.05 % FA) and eluent B (80 % ACN, 0.04 % FA) with a flow rate of 0.3 µL/min. The peptides were first desalted for 4 min at 5 % B, then separated with a gradient to 50 % B over 65 min, to 90 % B in 3 min, held at 90 % B for 8 min, and then equilibrated for 8 min at 4 % B.

For targeted absolute quantification, full MS spectra measurements were followed by parallel reaction monitoring (PRM) of targeted heavy AQUA peptides and the corresponding light, native peptide of the proteins of interest. The full MS spectra were acquired in profile mode with the following settings: resolution of 70000, mass range of 350–950 m/z, AGC target of 10^6^, and 80 ms maximum injection time. PRM MS2 spectra were acquired with an isolation window of 1.6 m/z, 17500 resolution, AGC target of 10^5^, 80 ms injection time, and an NCE of 27. The inclusion list contained 26 entries covering the light and heavy species of 10 peptides at the most intense charge state (2+ or 3+).

Database searches were performed with Proteome Discoverer (version 3.01.27; ThermoFisher). The raw data were compared with the UniProt protein databases of *S. cerevisiae* strain CEN.PK113-7D (UniProt 02.2023; 5,439 entries), and common contaminants using Sequest HT and Chimerys search algorithms. The Sequest search parameters were a semi-tryptic protease specificity with a maximum of 2 missed cleavage sites. The precursor mass tolerance was 8 ppm and the fragment mass tolerance was 0.02 Da. Oxidation of methionine and acetylation of protein N-termini were allowed as dynamic modifications. Carbamidomethylation of cysteine was set as a static modification. Chimerys database searches were performed with default settings. Percolator q-values were used to restrict the false discovery rate (FDR) of peptide spectrum matches to 0.01. The FDR of peptide and protein identifications was restricted to 1 % and strict parsimony principles were applied to protein grouping.

A spectral library of the AQUA peptides was generated using Proteome Discoverer with a sample containing only 500 fmol of the ten AQUA peptides, in addition to the PRM analyses of the spiked yeast samples. The PRM data of three concentrations analyzed in triplicate were imported and all transitions were reviewed using Skyline (57). Between four and nine transitions without interference were chosen for each peptide and the ratio of light and heavy peptides of the sum of transitions was calculated for absolute quantification. The list of used peptides and all corresponding transitions are provided in Table EV8.

### Calculation of absolute concentrations

Protein and metabolite concentrations are expressed as absolute intracellular concentrations. Metabolite concentrations initially expressed in µmol/g DCW were converted into absolute intracellular concentrations using a conversion factor of 6.59×10^10^ cells/g DCW (Fig EV4) and a cellular volume of 66 µm^3^ (58). The intracellular concentrations of PaCrtB and XdCrtI were calculated using a conversion factor of 0.63 g protein/g DCW for yeast grown on ammonium sulfate medium (59). All calculations for data conversion are provided in the corresponding Extended view Tables.

### Modeling

To test the impact of enzyme expression level on the obtained kinetic profiles, we built a toy kinetic model of the pathway under study. This model contains two metabolites (GGPP and phytoene) and three reactions: GGPP formation by CrtE, defined as a constant flux; GGPP conversion into phytoene by CrtB, modelled using a Michaelis-Menten rate law; and phytoene dilution by growth, modelled using mass action. The K_M_ value of CrtB was set arbitrarily to 1, and we simulated the steady-state phytoene production flux and GGPP concentration for three CrtB levels (V_max_ set to 0.02, 0.4 and 0.7) under a broad range of GGPP-producing flux (from 0.01 to 0.4 µM/h). The model has been developed with COPASI v4.39 (60) and is available from our GitHub repository (https://github.com/MetaSys-LISBP/in_vivo_enzymatic_parameters) and from the Biomodels database (61) under accession ID MODE<u>L2407240001</u>. All simulations were performed with COPASI.

### Calculation of enzyme parameters

Enzymatic parameters (V_max_ and K_M_) were determined by fitting a Michaelis-Menten equation to the measured relationships between substrate concentrations and reaction rates. Uncertainties on fitted parameters were determined using a Monte-Carlo approach. Briefly, 1000 simulated noisy datasets were generated (where the noise was determined as the standard deviation of residuals obtained for the fit of the experimental dataset), and the mean value, standard deviation, and 95 % confidence intervals of each parameter were determined from the distribution of values obtained for the 1000 datasets. For each enzyme, we verified that all parameters were identifiable based on the covariance matrix and on the results of the Monte-Carlo analysis. The code for parameter estimation and statistical analysis is provided as a Jupyter notebook at https://github.com/MetaSys-LISBP/in_vivo_enzymatic_parameters.

## Data Availability

The data used to make the figures can be found in the Source Data files. The model and computer code produced in this study are available in the following databases:

- Model: BioModels MODEL<u>2407240001</u>(https://www.ebi.ac.uk/biomodels/MODEL2407240001)
- Code: GitHub (https://github.com/MetaSys-LISBP/in_vivo_enzymatic_parameters)

## Supporting information

SourceData

## Acknowledgements

The authors thank MetaboHub-MetaToul (Metabolomics & Fluxomics facilities, Toulouse, France, https://mth-metatoul.com), which is part of the French National Infrastructure for Metabolomics and Fluxomics (https://www.metabohub.fr), funded by the ANR (MetaboHUB-ANR-11-INBS-0010), for access to MS facilities. The authors also thank Jean-Charles Portais (RESTORE, Geroscience & Rejuvenation Center, Université de Toulouse, INSERM, CNRS, EFS, Toulouse, France), Sylvie Dequin (SPO, Université Montpellier, INRAE, Institut Agro Montpellier, Montpellier, France), Thibault Nidelet (SPO, Université Montpellier, INRAE, Institut Agro Montpellier, Montpellier, France), Thomas Lautier (TBI, Université de Toulouse, CNRS, INRAE, INSA, Toulouse, France) and Sergueï Sokol (TBI, Université de Toulouse, CNRS, INRAE, INSA, Toulouse, France) for insightful discussions.

## Contributions

Funding acquisition: CC, SH, PM, GT; Project administration: CC, SH, GT; Conceptualization: CC, SCC, SH, PM, GT; Investigation: SCC, AC, HK, CT, AT, VG, CC, PM; Methodology: SCC, AC, HB, CT, FB, AT, VG, CC, SH, LGA, PM, GT; Software: PM; Formal Analysis: SCC, AC, HB, CT, AT, LGA, PM, GT; Visualization: SCC, PM, GT; Writing – original draft: SCC, PM, GT; Writing – review & editing: SCC, AC, HB, CT, FB, AT, VG, CC, SH, LGA, PM, GT; Supervision: SCC, VG, PM, GT.

## Expanded View Figure legends

Figure EV1. PaCrtB-FLAG expression determined by western blot in yeast strains with different GGPP concentrations (panels A, B and C correspond to three different biological replicates).

Figure EV2. Lycopene production in *S. cerevisiae*. A *S. cerevisiae* yGGPP34 strain expressing PDC1p-PaCrtB and TDH3p-BtCrtI. Bright field images were acquired using the camera LEICA DFC300FX mounted in the microscope Leica DM4000B with Leica EL6000 light source. Lycopene crystals are observed in red. B Decrease of specific growth rate in yeast strains expressing BtCrtI with different phytoene concentrations.

Figure EV3. Expression of the three studied CrtI-V5 proteins in strains with low (yGGPP030) and high (yGGPP044) phytoene content (panels A and B correspond to three different biological replicates).

Figure EV4. Correlation between cell/mL and OD_600nm_ (A) and between OD_600nm_ and mg DCW (B).

## Expanded View Table legends

Table EV1. Yeast strains used in this study.

Table EV2. PaCrtB and XdCrtI absolute protein quantification.

Table EV3. Enzymatic parameters measured *in vivo*.

Table EV4. *in vitro* parameters of bacterial phytoene synthase and phytoene desaturase.

Table EV5. Molecule number of each component for the *in vivo* enzymology reaction.

Table EV6. Plasmids used in this study.

Table EV7. Primers used for colony PCRs.

Table EV8. Peptides and PRM transitions used for absolute quantification of the enzymes.

## References

1. S. R. McGuffee, A. H. Elcock, Diffusion, Crowding & Protein Stability in a Dynamic Molecular Model of the Bacterial Cytoplasm. PLOS Computational Biology 6, e1000694 (2010).

2. B. M. B. Karen van Eunen, The importance and challenges of *in vivo*-like enzyme kinetics. Perspectives in Science 1, 126–130 (2014).

3. D. Ringe, G. A. Petsko, How Enzymes Work. Science 320, 1428–1429 (2008).

4. A. Sancar, Structure and Function of Photolyase and *in vivo* Enzymology: 50th Anniversary. J. Biol. Chem. 283, 32153–32157 (2008).

5. W. Harm, H. Harm, C. S. Rupert, Analysis of photoenzymatic repair of UV lesions in DNA by single light flashes: II. in vivo studies with Escherichia coli cells and bacteriophage. Mutation Research/Fundamental and Molecular Mechanisms of Mutagenesis 6, 371–385 (1968).

6. B. E. Wright, M. H. Butler, K. R. Albe, Systems analysis of the tricarboxylic acid cycle in Dictyostelium discoideum. I. The basis for model construction. Journal of Biological Chemistry 267, 3101–3105 (1992).

7. D. Heckmann, et al., Kinetic profiling of metabolic specialists demonstrates stability and consistency of *in vivo* enzyme turnover numbers. Proc Natl Acad Sci USA 202001562 (2020). 10.1073/pnas.2001562117.

8. D. Davidi, et al., Global characterization of *in vivo* enzyme catalytic rates and their correspondence to *in vitro* k _cat_ measurements. Proc Natl Acad Sci USA 113, 3401–3406 (2016).

9. A. Zotter, F. Bäuerle, D. Dey, V. Kiss, G. Schreiber, Quantifying enzyme activity in living cells. J Biol Chem 292, 15838–15848 (2017).

10. S. Bhattacharyya, S. Bershtein, B. V. Adkar, J. Woodard, E. I. Shakhnovich, Metabolic response to point mutations reveals principles of modulation of *in vivo* enzyme activity and phenotype. Mol Syst Biol 17, e10200 (2021).

11. Y. Takakuwa, H. Nishino, Y. Ishibe, T. Ishibashi, Properties and kinetics of membrane-bound enzymes when both the enzyme and substrate are components of the same microsomal membrane. Studies on lathosterol 5-desaturase. Journal of Biological Chemistry 269, 27889– 27893 (1994).

12. M. Camagna, et al., Enzyme Fusion Removes Competition for Geranylgeranyl Diphosphate in Carotenogenesis. Plant Physiol. 179, 1013–1027 (2019).

13. P. Urban, et al., Engineered yeasts simulating P450-dependent metabolisms: tricks, myths and reality. Drug Metabol Drug Interact 11, 169–200 (1994).

14. S. Gemmecker, et al., Phytoene Desaturase from Oryza sativa: Oligomeric Assembly, Membrane Association and Preliminary 3D-Analysis. PLoS One 10, e0131717 (2015).

15. D. Iwata-Reuyl, S. K. Math, S. B. Desai, C. D. Poulter, Bacterial Phytoene Synthase:1 Molecular Cloning, Expression, and Characterization of Erwinia herbicola Phytoene Synthase†. Biochemistry 42, 3359–3365 (2003).

16. P. Schaub, et al., On the Structure and Function of the Phytoene Desaturase CRTI from Pantoea ananatis, a Membrane-Peripheral and FAD-Dependent Oxidase/Isomerase. PLoS ONE 7, e39550 (2012).

17. M. Fournié, G. Truan, Multiplicity of carotene patterns derives from competition between phytoene desaturase diversification and biological environments. Sci Rep 10, 21106 (2020).

18. P. Stickforth, G. Sandmann, Structural and kinetics properties of a mutated phytoene desaturase from Rubrivivax gelatinosus with modified product specificity. Arch Biochem Biophys 505, 118–122 (2011).

19. L. Michaelis, M. Menten, Die Kinetik der Invertinwirkung. Biochemische Zeitschrift 49, 333–369 (1913).

20. F. J. Bruggeman, J. J. Hornberg, F. C. Boogerd, H. V. Westerhoff, Introduction to systems biology. EXS 97, 1–19 (2007).

21. U. Neudert, I. M. Martinez-Férez, P. D. Fraser, G. Sandmann, Expression of an active phytoene synthase from Erwinia uredovora and biochemical properties of the enzyme. Biochimica et Biophysica Acta (BBA) - Lipids and Lipid Metabolism 1392, 51–58 (1998).

22. P. D. Fraser, W. Schuch, P. M. Bramley, Phytoene synthase from tomato (Lycopersicon esculentum) chloroplasts – partial purification and biochemical properties. Planta 211, 361–369 (2000).

23. B. Camara, “[32] Plant phytoene synthase complex: Component enzymes, immunology, and biogenesis” in Methods in Enzymology, Carotenoids Part B: Metabolism, Genetics, and Biosynthesis., (Academic Press, 1993), pp. 352–365.

24. A. Schofield, G. Paliyath, Modulation of carotenoid biosynthesis during tomato fruit ripening through phytochrome regulation of phytoene synthase activity. Plant Physiology and Biochemistry 43, 1052–1060 (2005).

25. G. Sandmann, Evolution of carotene desaturation: The complication of a simple pathway. Archives of Biochemistry and Biophysics 483, 169–174 (2009).

26. J. Koschmieder, et al., Plant-type phytoene desaturase: Functional evaluation of structural implications. PLoS ONE 12, e0187628 (2017).

27. H. Rabeharindranto, et al., Enzyme-fusion strategies for redirecting and improving carotenoid synthesis in *S. cerevisiae*. Metab Eng Commun 8, e00086 (2019).

28. Y. Chen, et al., Lycopene overproduction in *Saccharomyces cerevisiae* through combining pathway engineering with host engineering. Microbial Cell Factories 15, 113 (2016).

29. P. J. Westfall, et al., Production of amorphadiene in yeast, and its conversion to dihydroartemisinic acid, precursor to the antimalarial agent artemisinin. PNAS 109, E111–E118 (2012).

30. J. Zhao, et al., Dynamic control of *ERG20* expression combined with minimized endogenous downstream metabolism contributes to the improvement of geraniol production in *Saccharomyces cerevisiae*. Microb Cell Fact 16, 1–11 (2017).

31. A. Faulkner, et al., The LPP1 and *DPP1* Gene Products Account for Most of the Isoprenoid Phosphate Phosphatase Activities in*Saccharomyces cerevisiae*. J. Biol. Chem. 274, 14831–14837 (1999).

32. B. Peng, L. K. Nielsen, S. C. Kampranis, C. E. Vickers, Engineered protein degradation of farnesyl pyrophosphate synthase is an effective regulatory mechanism to increase monoterpene production in *Saccharomyces cerevisiae*. Metabolic Engineering 47, 83–93 (2018).

33. C. Chambon, V. Ladeveze, A. Oulmouden, M. Servouse, F. Karst, Isolation and properties of yeast mutants affected in farnesyl diphosphate synthetase. Curr Genet 18, 41–46 (1990).

34. K. Tokuhiro, et al., Overproduction of Geranylgeraniol by Metabolically Engineered *Saccharomyces cerevisiae*. Appl. Environ. Microbiol. 75, 5536–5543 (2009).

35. X. Zhou, et al., Phytoene Synthase: The Key Rate-Limiting Enzyme of Carotenoid Biosynthesis in Plants. Frontiers in Plant Science 13 (2022).

36. D. Leys, J. Basran, F. Talfournier, M. J. Sutcliffe, N. S. Scrutton, Extensive conformational sampling in a ternary electron transfer complex. Nat Struct Mol Biol 10, 219–225 (2003).

37. 37. L. Phégnon, et al., Phosphogluconolactonase as the linchpin of an efficient pentose phosphate pathway. [Preprint] (2023). Available at: https://www.biorxiv.org/content/10.1101/2023.11.29.569206v2 [Accessed 8 December 2023].

38. J. O. Park, et al., Metabolite concentrations, fluxes and free energies imply efficient enzyme usage. Nat Chem Biol 12, 482–489 (2016).

39. X. Bu, J. Lin, C. Duan, M. A. G. Koffas, G. Yan, Dual regulation of lipid droplet-triacylglycerol metabolism and ERG9 expression for improved β-carotene production in *Saccharomyces cerevisiae*. Microb Cell Fact 21, 3 (2022).

40. Y. Zhu, et al., Enhanced synthesis of Coenzyme Q10 by reducing the competitive production of carotenoids in Rhodobacter sphaeroides. Biochemical Engineering Journal 125, 50–55 (2017).

41. R. Heinrich, T. A. Rapoport, A linear steady-state treatment of enzymatic chains. General properties, control and effector strength. Eur J Biochem 42, 89–95 (1974).

42. H. Kacser, J. A. Burns, The control of flux. Symp Soc Exp Biol 27, 65–104 (1973).

43. B. Enjalbert, P. Millard, M. Dinclaux, J.-C. Portais, F. Létisse, Acetate fluxes in Escherichia coli are determined by the thermodynamic control of the Pta-AckA pathway. Sci Rep 7, 42135 (2017).

44. S.-M. Fendt, et al., Tradeoff between enzyme and metabolite efficiency maintains metabolic homeostasis upon perturbations in enzyme capacity. Mol Syst Biol 6, 356 (2010).

45. P. Millard, K. Smallbone, P. Mendes, Metabolic regulation is sufficient for global and robust coordination of glucose uptake, catabolism, energy production and growth in Escherichia coli. PLOS Computational Biology 13, e1005396 (2017).

46. M. L. Reaves, B. D. Young, A. M. Hosios, Y.-F. Xu, J. D. Rabinowitz, Pyrimidine homeostasis is accomplished by directed overflow metabolism. Nature 500, 237–241 (2013).

47. R. Verwaal, et al., High-Level Production of Beta-Carotene in *Saccharomyces cerevisiae* by Successive Transformation with Carotenogenic Genes from Xanthophyllomyces dendrorhous. Appl. Environ. Microbiol. 73, 4342–4350 (2007).

48. F. X. Cunningham Jr, et al., Functional analysis of the beta and epsilon lycopene cyclase enzymes of Arabidopsis reveals a mechanism for control of cyclic carotenoid formation. The Plant Cell 8, 1613–1626 (1996).

49. W. Xie, et al., Construction of a controllable β-carotene biosynthetic pathway by decentralized assembly strategy in *Saccharomyces cerevisiae*. Biotechnol. Bioeng. 111, 125–133 (2014).

50. Engineering of Promoter Replacement Cassettes for Fine-Tuning of Gene Expression in *Saccharomyces cerevisiae* | Applied and Environmental Microbiology. Available at: https://journals-asm-org.insb.bib.cnrs.fr/doi/10.1128/aem.00530-06 [Accessed 30 January 2024].

51. R. D. Gietz, Yeast transformation by the LiAc/SS carrier DNA/PEG method. Methods Mol. Biol. 1205, 1–12 (2014).

52. U. Güldener, S. Heck, T. Fiedler, J. Beinhauer, J. H. Hegemann, A New Efficient Gene Disruption Cassette for Repeated Use in Budding Yeast. Nucl. Acids Res. 24, 2519–2524 (1996).

53. C. Verduyn, E. Postma, W. A. Scheffers, J. P. Y. 1990 van Dijken, Physiology of *Saccharomyces cerevisiae* in Anaerobic Glucose-Limited Chemostat Culturesx. Microbiology 136, 395–403.

54. P. Millard, et al., ScalaFlux: A scalable approach to quantify fluxes in metabolic subnetworks. PLOS Computational Biology 16, e1007799 (2020).

55. S. Castaño-Cerezo, et al., Functional analysis of isoprenoid precursors biosynthesis by quantitative metabolomics and isotopologue profiling. Metabolomics 15, 115 (2019).

56. T. Zhang, et al., An improved method for whole protein extraction from yeast *Saccharomyces cerevisiae*. Yeast 28, 795–798 (2011).

57. B. MacLean, et al., Effect of Collision Energy Optimization on the Measurement of Peptides by Selected Reaction Monitoring (SRM) Mass Spectrometry. Anal. Chem. 82, 10116–10124 (2010).

58. N. Punekar, “In Vitro Versus In Vivo: Concepts and Consequences” in ENZYMES: Catalysis, Kinetics and Mechanisms, Springer Singapore, (2018), pp. 493–519.

59. E. Albers, C. Larsson, G. Lidén, C. Niklasson, L. Gustafsson, Influence of the nitrogen source on *Saccharomyces cerevisiae* anaerobic growth and product formation. Appl Environ Microbiol 62, 3187–3195 (1996).

60. COPASI—a COmplex PAthway SImulator | Bioinformatics | Oxford Academic. Available at: https://academic.oup.com/bioinformatics/article/22/24/3067/208398 [Accessed 25 July 2024].

61. R. S. Malik-Sheriff, et al., BioModels—15 years of sharing computational models in life science. Nucleic Acids Research 48, D407–D415 (2020).

