## Supplementary figures and images for "Combining systems and synthetic biology for in vivo enzymology"

### EV1A_CROP.TIF

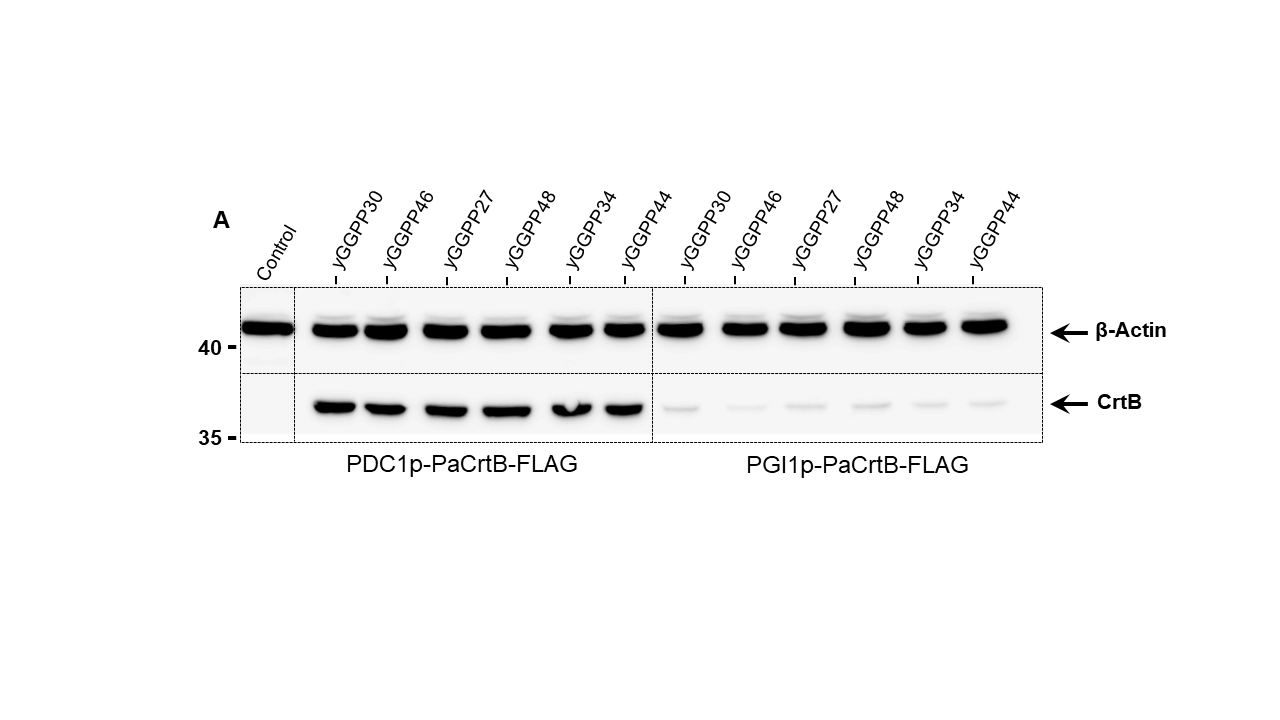

### EV1A_FULL.TIF

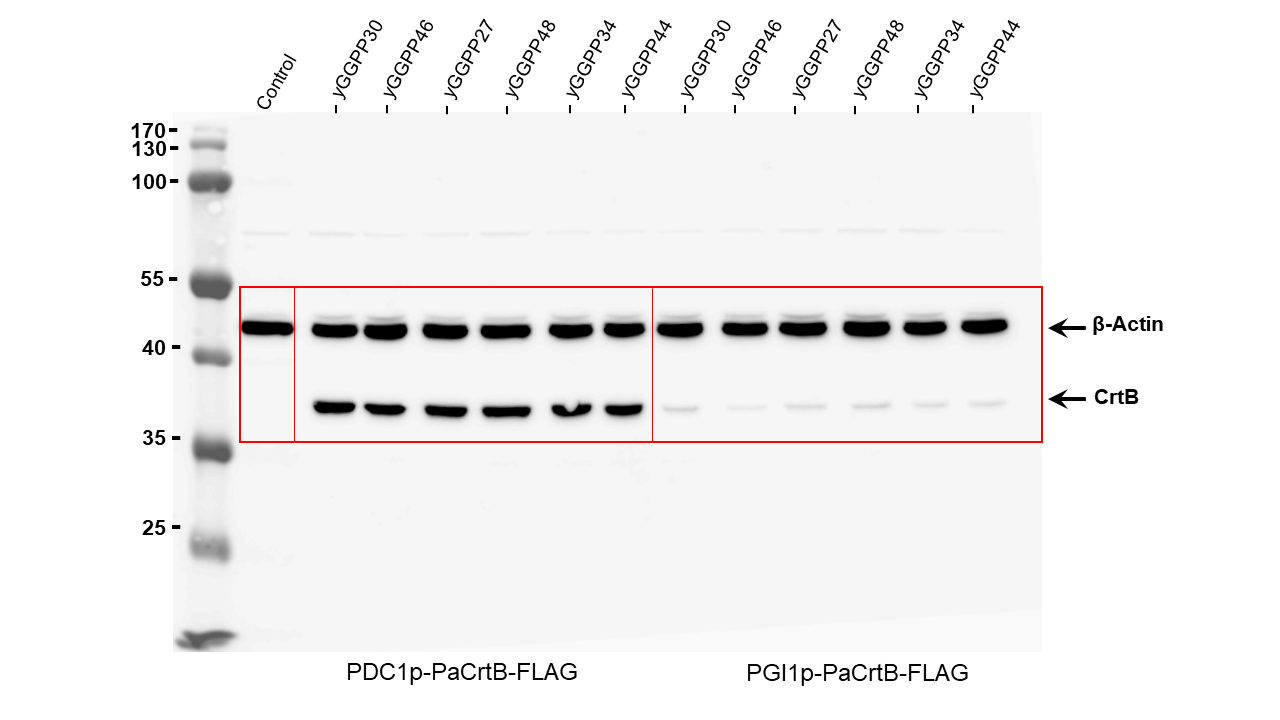

### EV1B_CROP.TIF

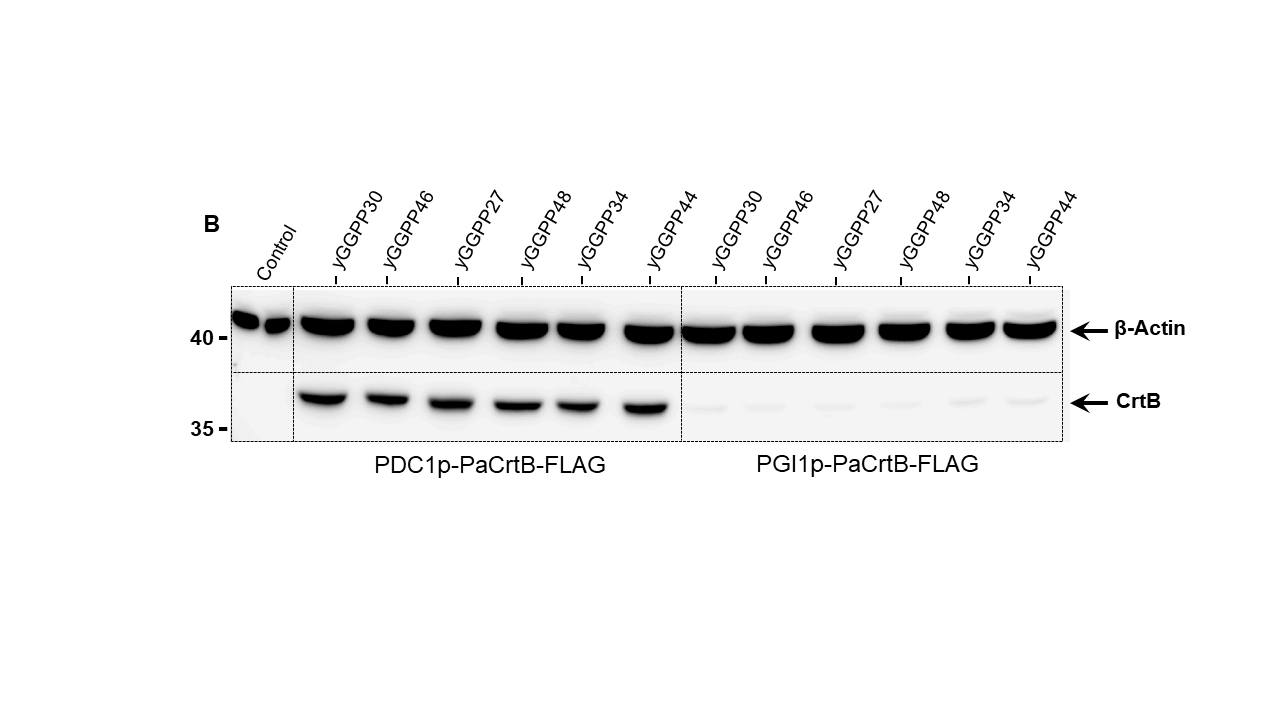

### EV1B_FULL.TIF

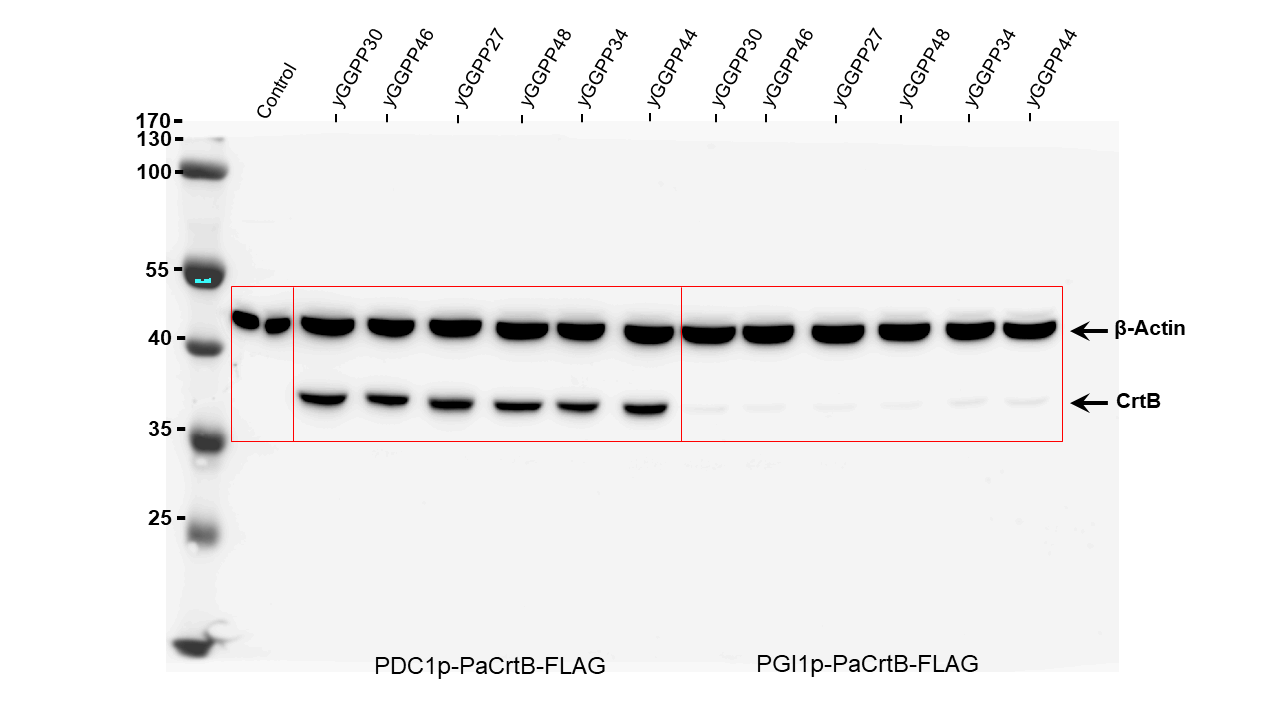

### EV1C_CROP.TIF

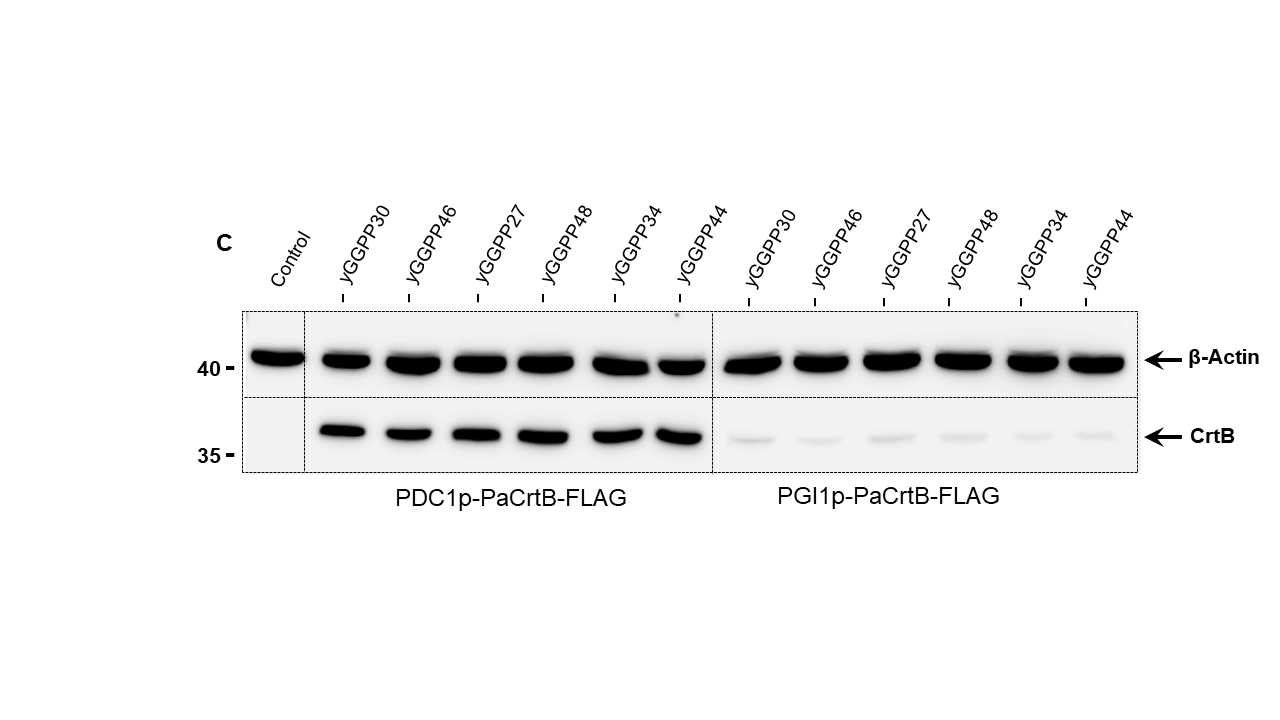

### EV1C_FULL.TIF

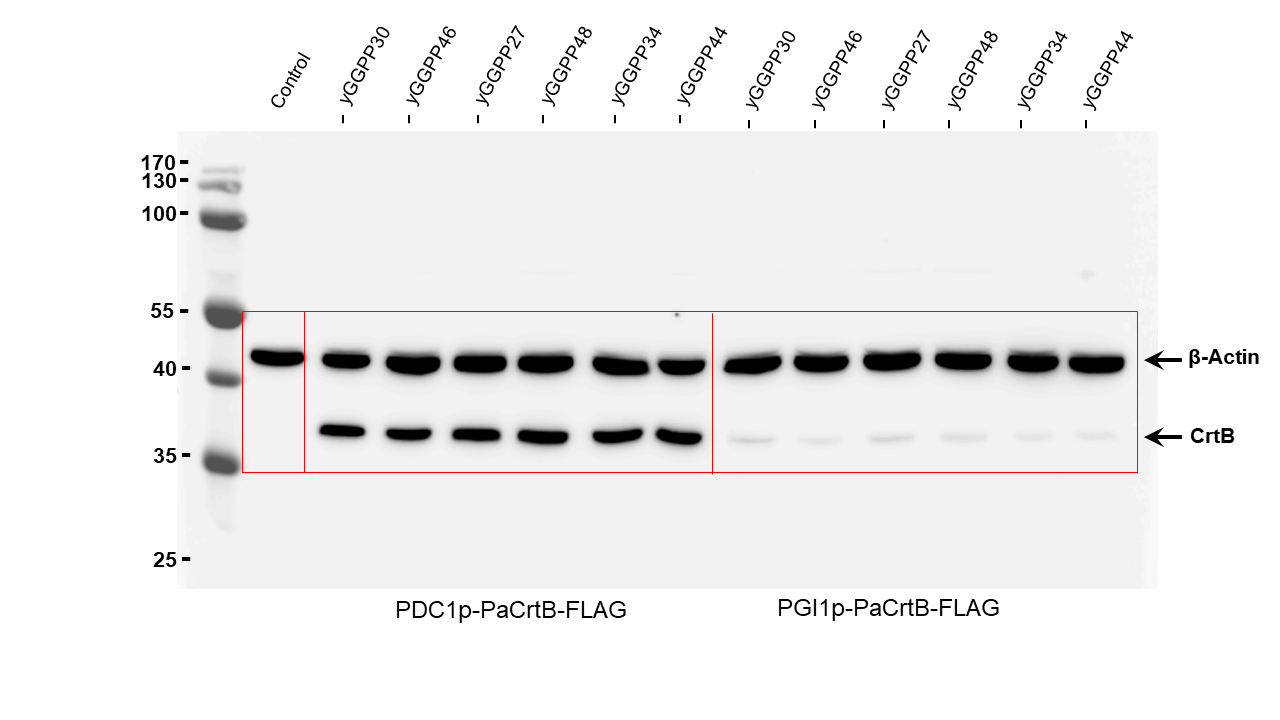

### EV2A.tif

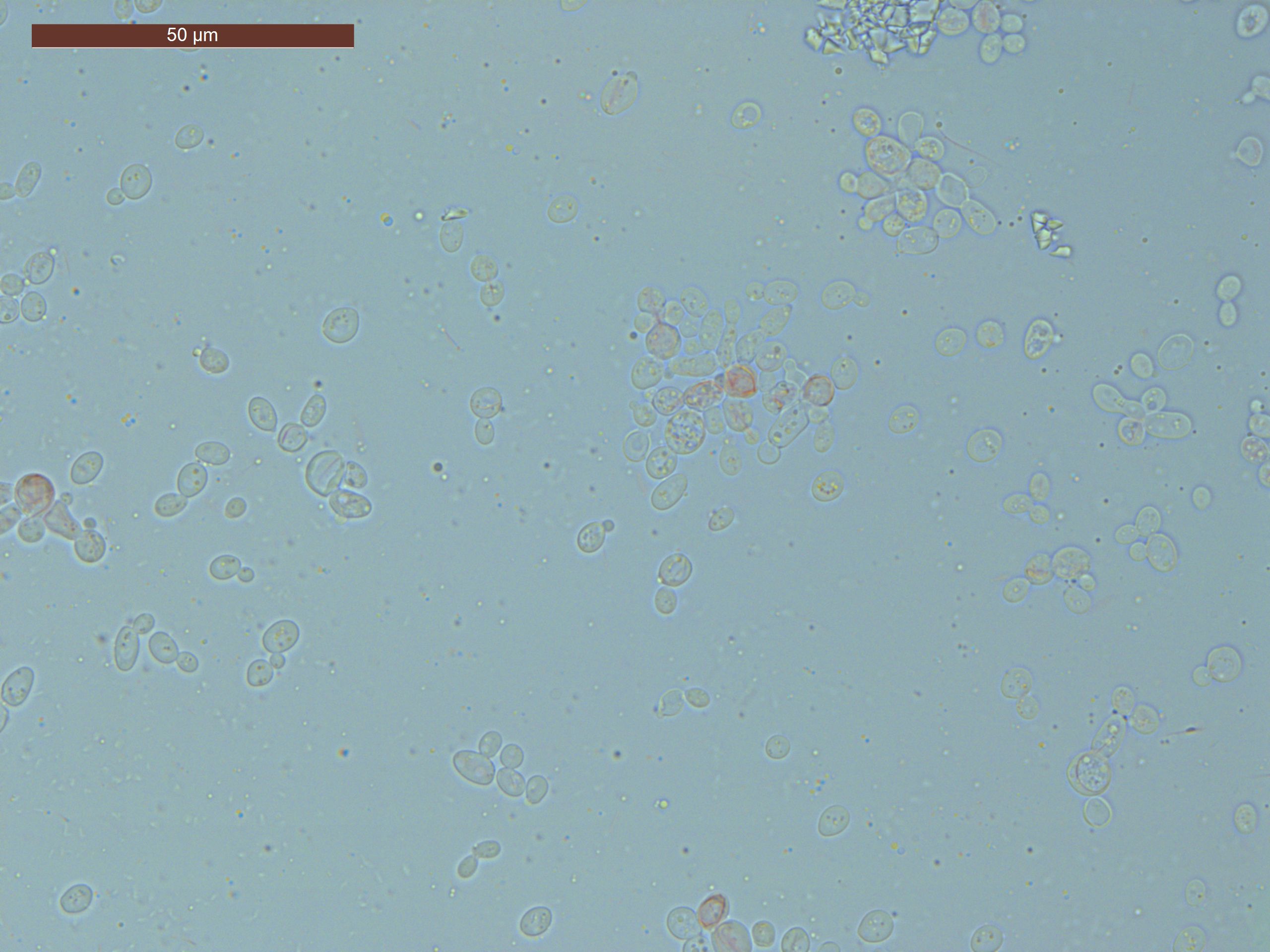

### EV3A_CROP.TIF

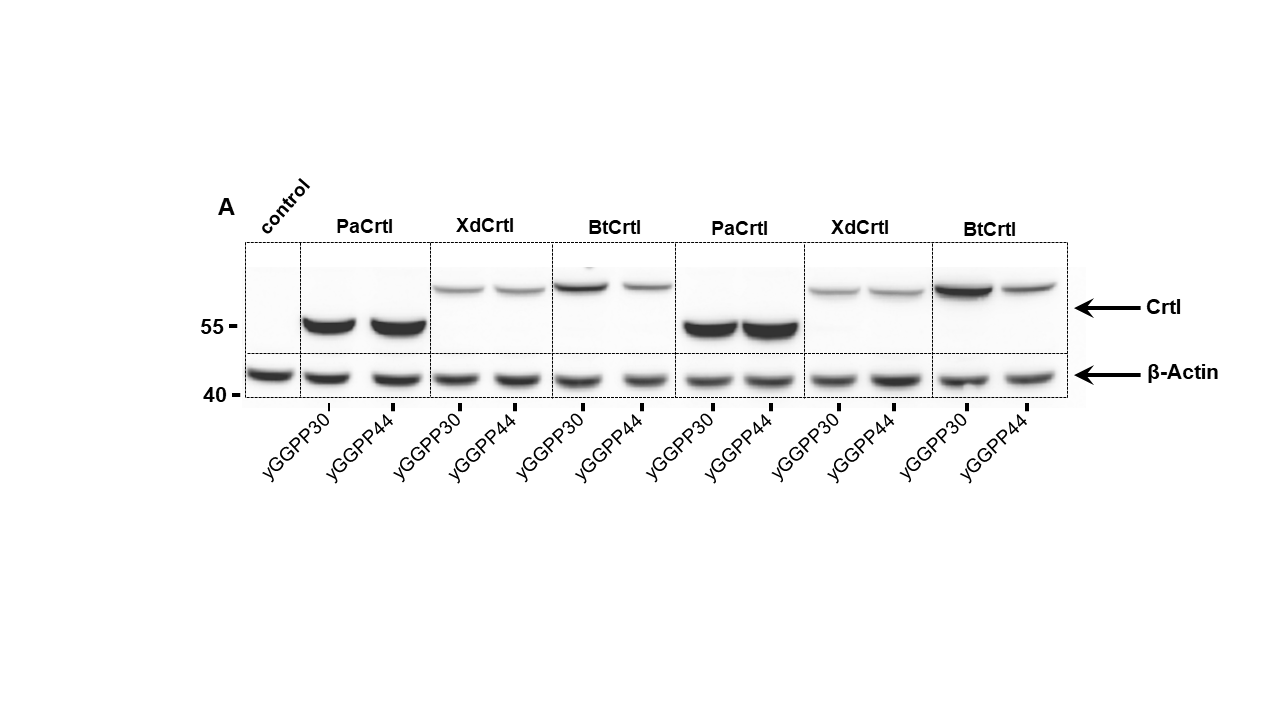

### EV3A_FULL.TIF

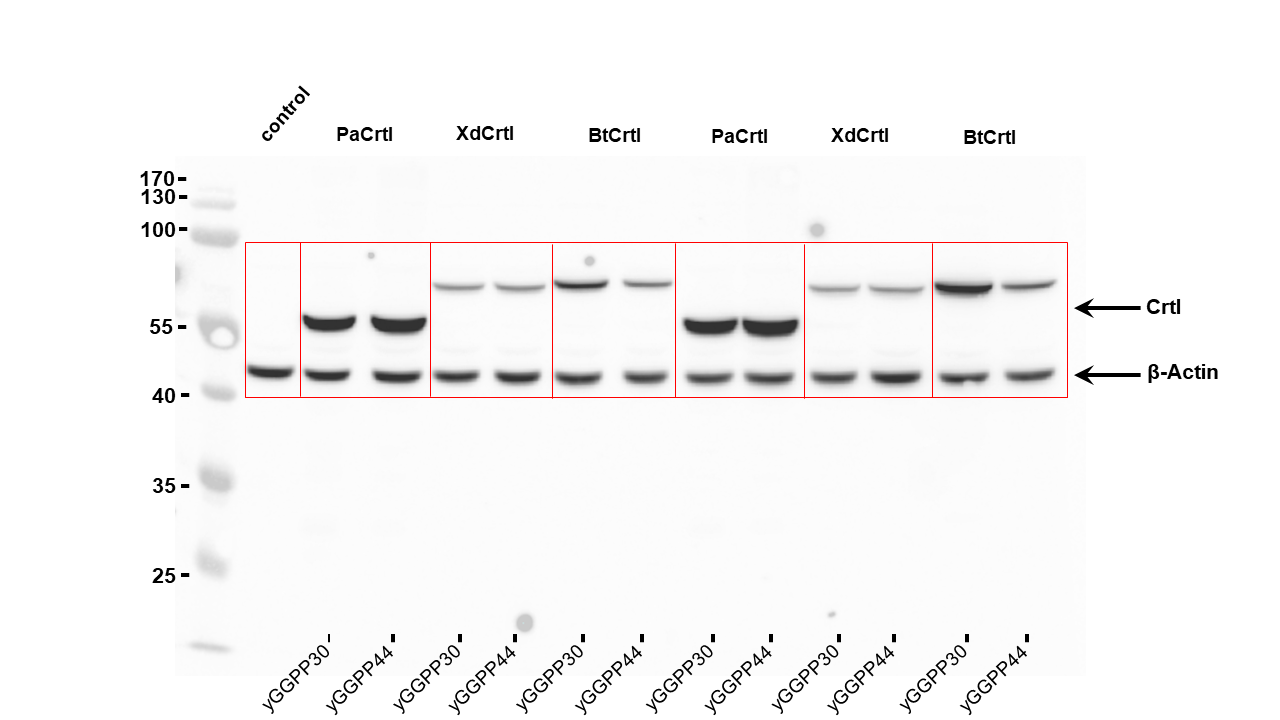

### EV3B_CROP.TIF

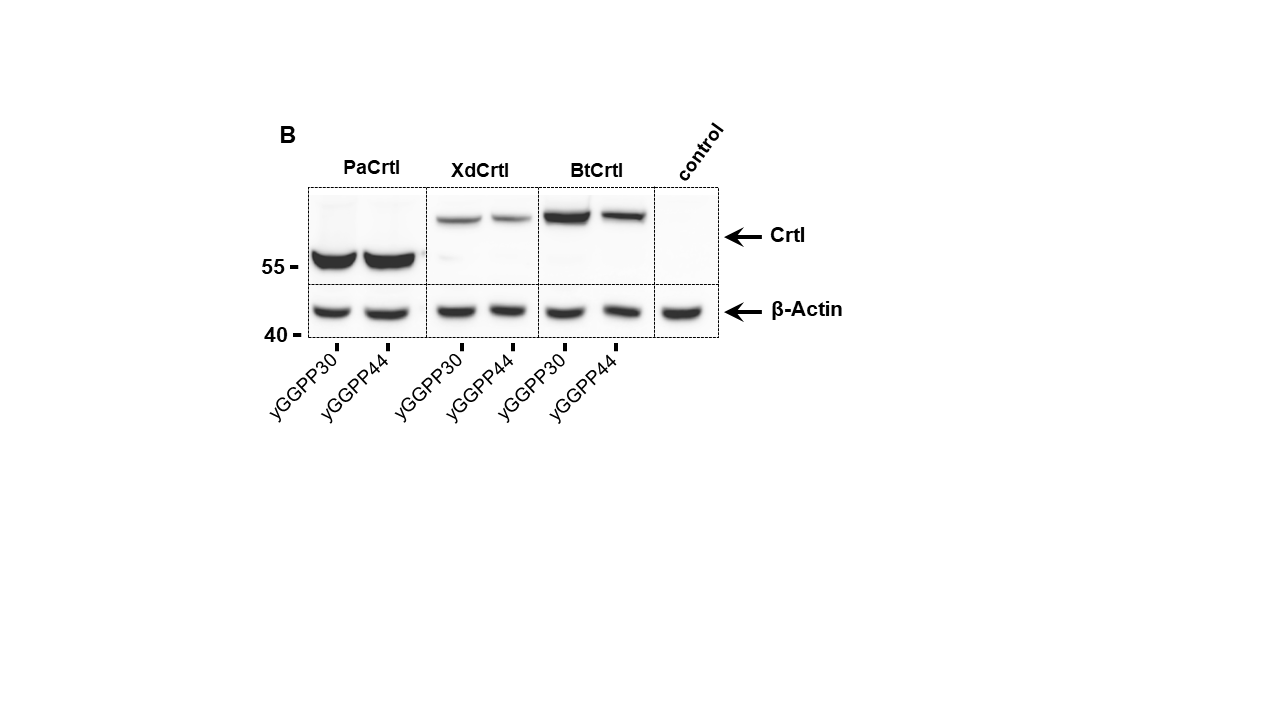

### EV3B_FULL.TIF

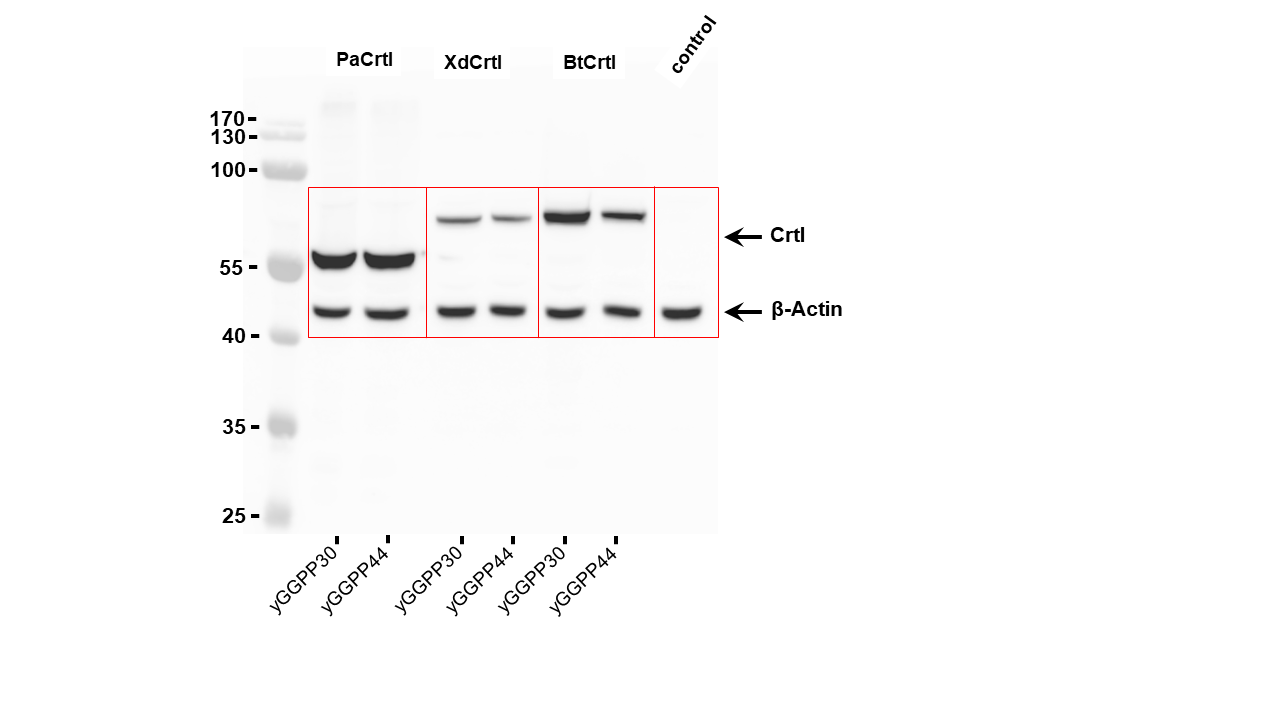

### Figure EV1

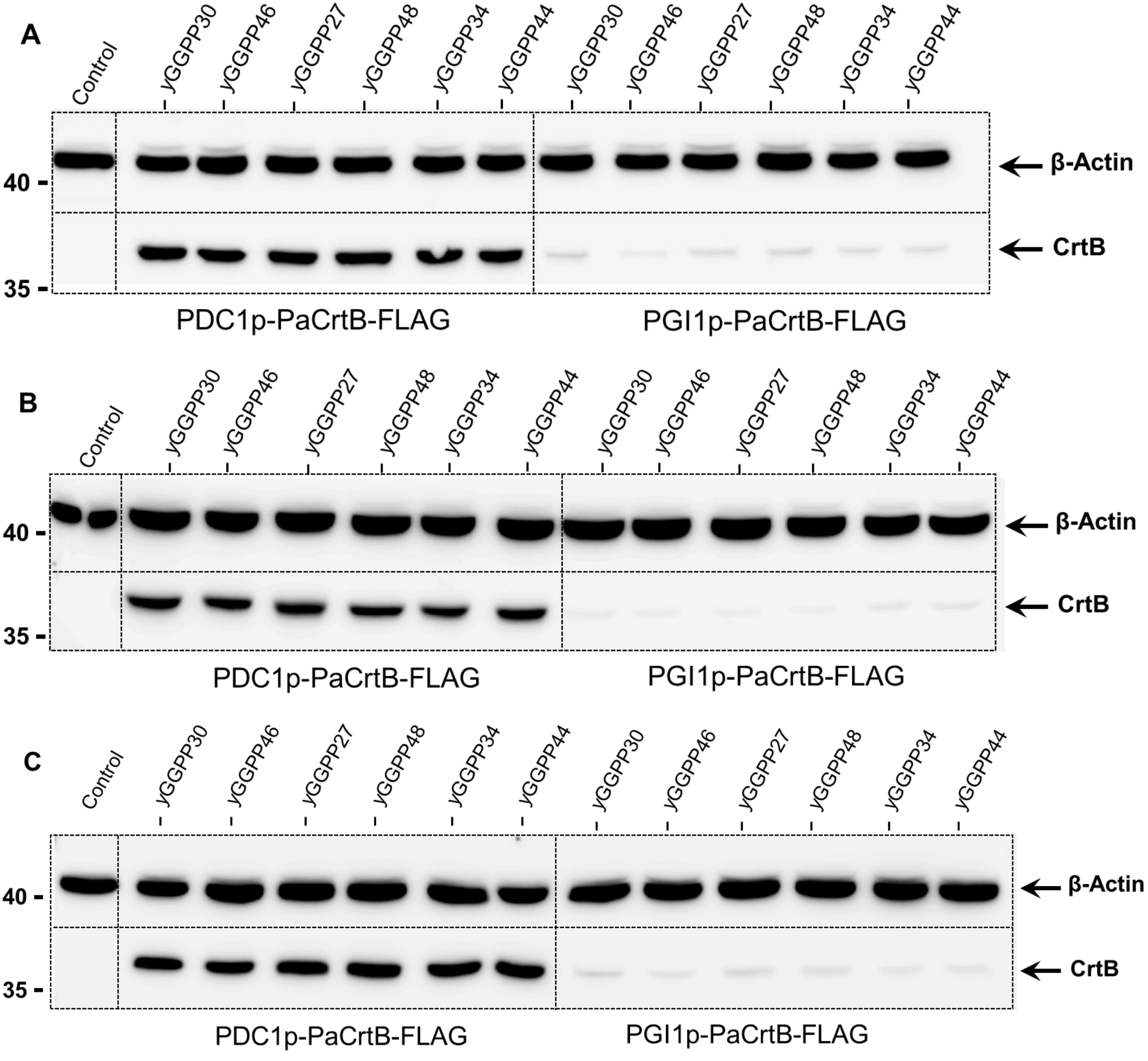

### Figure EV2

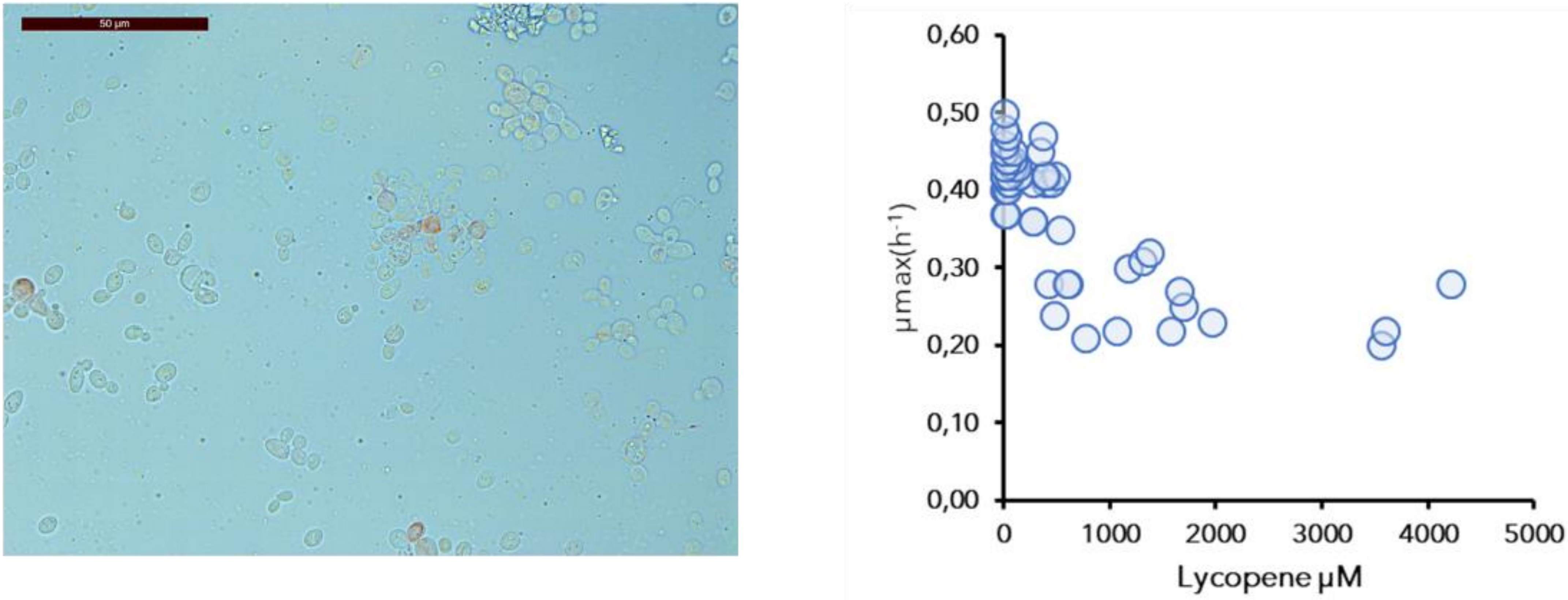

### Figure EV3

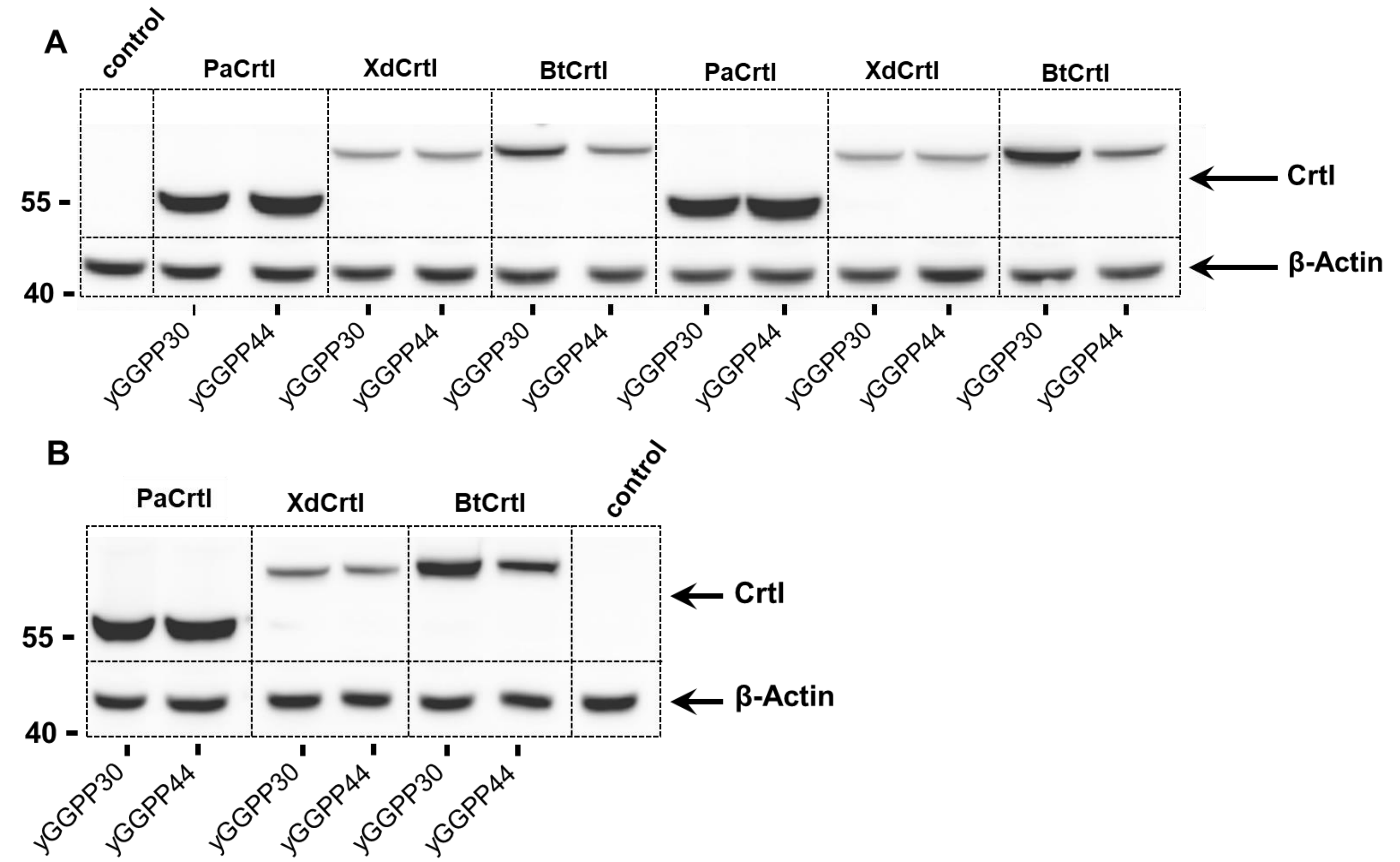

### Figure EV4

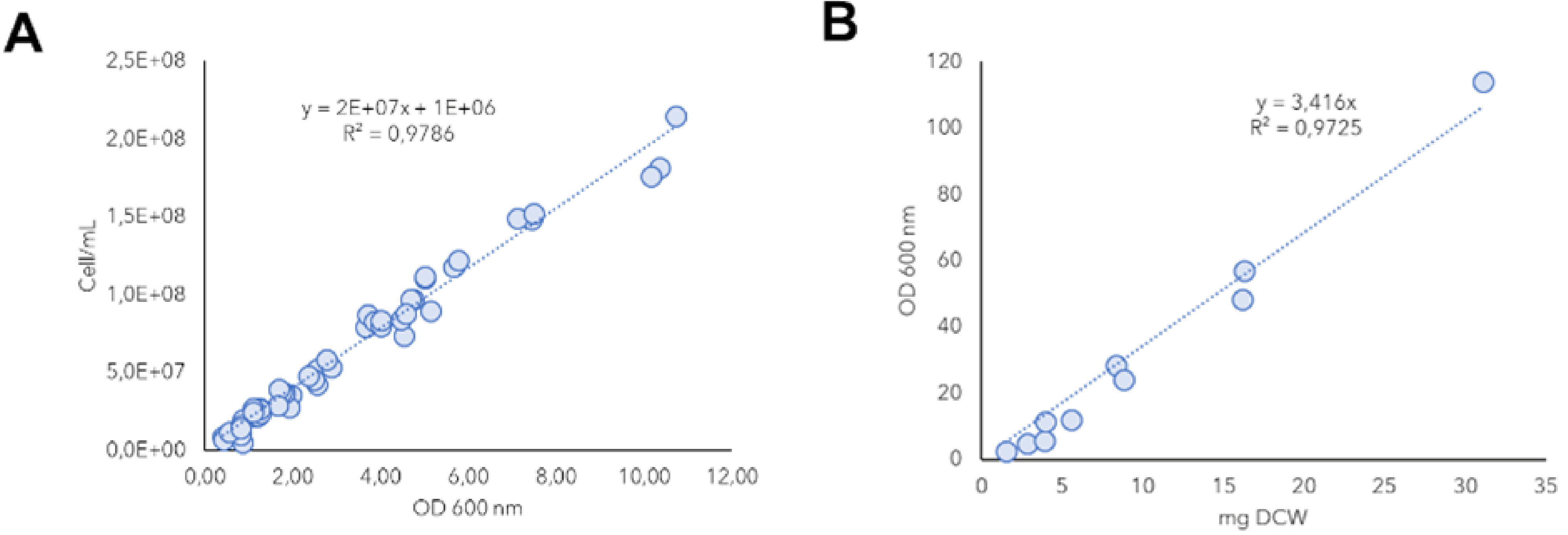
